# Arabidopsis inositol polyphosphate kinases IPK1 and ITPK1 modulate crosstalks between SA-dependent immunity and phosphate-starvation responses

**DOI:** 10.1101/2021.01.27.428180

**Authors:** Hitika Gulabani, Krishnendu Goswami, Yashika Walia, Jewel Jameeta Noor, Kishor D. Ingole, Abhisha Roy, Debabrata Laha, Gabriel Schaaf, Saikat Bhattacharjee

## Abstract

The propensity for polyphosphorylation makes *myo*-inositol derivatives, the inositol polyphosphates (InsPs), especially phytic acid or inositol hexakisphosphate (InsP_6_) the major form of phosphate storage in plants. Acts of pyrophosphorylation on InsP_6_ generates InsP_7_ or InsP_8_ containing high-energy phosphoanhydride bonds that are harnessed during energy requirements of a cell. Also implicated as co-factors for several phytohormone signaling networks, InsP_7_/InsP_8_ modulate key developmental processes. With recent identification as the common moeity for transducing both jasmonic acid (JA) and phosphate-starvation responses (PSR), InsP_8_ is the classic example of a metabolite that may moonlight crosstalks to different cellular pathways during diverse stress adaptations. We show here that *Arabidopsis thaliana* INOSITOL PENTAKISPHOSPHATE 2-KINASE (IPK1), INOSITOL 1,3,4-TRISPHOSPHATE 5/6-KINASE 1 (ITPK1), and DIPHOSPHOINOSITOL PENTAKISPHOSPHATE KINASE 2 (VIH2), but not other InsP-kinases, suppress basal salicylic acid (SA)-dependent immunity. In *ipk1, itpk1* or *vih2* mutants, elevated endogenous SA levels and constitutive activation of defense signaling lead to enhanced resistance against the virulent *Pseudomonas syringae pv* tomato DC3000 (*PstDC300*0) strain. Our data reveal that activated SA-signaling sectors in these mutants modulate expression amplitudes of phosphate-starvation inducible (PSI)-genes, reported earlier. In turn, via mutualism the heightened basal defenses in these mutants require upregulated PSI-gene expressions likely highlighting the increased demand of phosphates required to support immunity. We demonstrate that SA is induced in phosphate-deprived plants, however its defense-promoting functions are likely diverted to PSR-supportive roles. Overall, our investigations reveal selective InsPs as crosstalk mediators among diverse signaling networks programming stress-appropriate adaptations.

## Introduction

Several evidences illustrate the versatility and relevance of inositol (Ins) derivatives in regulating cellular homeostasis in plants (Majerus, 1992; Murthy, 1996; Gillaspy, 2011; Gillaspy, 2013). The availability of six hydroxyl groups on a cyclic alcohol and its propensity to attach varied number of phosphates generates diverse water soluble InsP moieties with roles in cellular transduction of signals in multiple pathways (Shears *et al*., 2012). Evidentially, tight regulations in maintaining cellular ratios of different InsPs are crucial and disturbed only spatiotemporally during cellular responses (Monserrate and York, 2010; Mishkind *et al*., 2009). As a potent chelator of polyvalent metal ions especially Ca^2+^, Mg^2+^, Zn^2+^, Fe^2+^ among others, although InsP_6_ is considered an anti-nutrient, recent studies identify anti-oxidative benefits of these roles in therapeutic interventions (Silva and Bracarense, 2016). The phospholipid conjugates phosphatidylinositols (PtdIns) though comprise only a minor pool of membrane lipids nevertheless is pivotal in determining cell architectures, facilitate protein anchors, regulate vesicle trafficking, organizes cytoskeleton, and also serve as sources for signaling InsPs via the lipid-dependent pathway (Munnik and Nielsen, 2011; Boss and Im, 2012). Dynamic changes in localized PtdIns are orchestrated by phosphatases and phospholipases and are key players in downstream signaling during regular development or responses to (a)biotic stresses (Dieck *et al*., 2012; Heilmann, 2016).

Glucose-6-phosphate, the precursor molecule for InsP biosynthesis is converted to *myo*-inositol-3-phosphate (Ins3P) by MYO-INOSITOL PHOSPHATE SYNTHASES (MIPS1-3 in Arabidopsis) (Donahue *et al*., 2010). Subsequent dephosphorylation of Ins3P by INOSITOL MONOPHOSPHATASE (IMP) generates *myo*-inositol (Ins) for conjugation with phospholipids via membrane-localized PHOSPHATIDYLINOSITOL SYNTHASE (PIS). Ins is phosphorylated by PtdIns4-KINASE (PI4K) and PtdIns4P 5-KINASES (PIP5K) for PtdIns4P and PtdIns(4,5)P_2_ synthesis. Biotic or abiotic stimuli cause activation of PHOSPHOLIPASE C (PLC) that hydrolyzes PtdIns4P or PtdIns(4,5)P_2_ producing diacylglycerol (DAG) and inositol trisphosphate (InsP_3_) that have important roles as signaling transducers of downstream responses (Gillaspy, 2011; Hong *et al*., 2016). InsP_3_ causes increase in intracellular calcium (Ca^2+^) via its liberation from endogenous stores although a *bona fide* receptor for InsP_3_-gated calcium release remains unidentified in plants (Berridge, 1993; Lee *et al*., 1996; Gillaspy, 2011; Boss and Im, 2012). Increasing studies indicate that instead of being the direct mediator of intracellular Ca^2+^ spike, in plants as well as in yeast, InsP_3_ is converted primarily to InsP_6_ for this function. In response to drought stress or ABA treatment, InsP_3_ as well as InsP_6_ levels in guard cells are elevated causing stomatal closure (Lee *et al*., 1996; Staxen *et al*., 1999; Lemtiri-Chlieh *et al*., 2003). Controlled release of InsP_3_ or InsP_6_ from their respective caged-conjugates coincides with mobilization of Ca^2+^ from endomembrane reserves (Lemtiri-Chlieh *et al*., 2003). Again, the identity of the receptor that facilitates InsP_6_-mediated Ca^2+^ spike remains elusive.

In the lipid-dependent origin of InsP_6_, sequential phosphorylation of InsP_3_ are performed by the redundant INOSITOL POLYPHOSPHATE 6-/3-KINASES (IPK2α and IPK2β) producing inositol tetrakisphosphate (InsP_4_) and inositol-pentakisphosphate (InsP_5_) (Xia *et al*., 2003; Stevenson-Paulik *et al*., 2005). An alternate lipid-independent pathway performing *de novo* synthesis of InsPs exists unique to plants and closely related protists (Stevenson-Paulik *et al*., 2005; Desfougeres *et al*., 2019). In this mode, via coordinated activities of myo-INOSITOL KINASE (MIK) and INOSITOL-TETRAKISPHOSPHATE 5,6-KINASES (ITPKs), InsP is sequentially phosphorylated to generate InsP_5_ (Shi *et al*., 2005). In Arabidopsis, a single gene-encoded INOSITOL PENTAKISPHOSPHATE 2-KINASE (IPK1) catalyzes the synthesis of InsP_6_ from InsP_5_ (Sweetman *et al*., 2007). An *IPK1* null-mutant is lethal and plants with partial loss-of-function (*ipk1-1*) display developmental defects indicating the importance of this abundant InsP (Kuo *et al*., 2014). Low InsP_6_ plants are categorized as *lpa* (low phytic acid) and include *ipk2β-1, mrp5, itpk1-2, itpk4-1*, and *mik-1* (Stevenson-Paulik *et al*., 2005; Kim and Tai, 2011). Anti-nutrient property of InsP_6_ in humans and monogastric animals along with its negative impact as an environmental pollutant due to eutrophication has instigated continuous agricultural efforts to develop *lpa* crops (Turner *et al*., 2002). However, with noted developmental consequences in several *lpa* mutants such endeavors are a constant concern to overall plant health (Lee *et al*., 2015; Donahue *et al*., 2010). InsP_6_ is a further substrate for pyrophosphorylation by ITPK1/2 to generate InsP_7_ (Adepoju *et al*., 2019; Laha *et al*., 2019) and further to InsP_8_ by VIH1/2 kinases (Desai *et al*., 2014; Laha *et al*., 2015). During the last decade, co-factor roles of specific plant InsPs have gained prominence. Demonstrated by structural and mutational studies, InsP_6_/InsP_7_ role as cofactor in auxin-responses and InsP_5_/InsP_8_ for jasmonic-acid signaling have been deciphered (Tan *et al*., 2007; Sheard *et al*., 2010; Laha *et al*., 2015; Laha *et al*., 2020). More recently, InsP_8_ is also implicated in maintenance of Pi-homeostasis and regulates PSR (phosphate-starvation response) upon Pi-deprivation (Kuo *et al*., 2014; Kuo *et al*., 2018; Ried *et al*., 2019; Zhu *et al*., 2019; Dong *et al*., 2019).

Incorporating such functional diversities in cellular processes, InsP implications on innate immune signaling of plants is undoubtedly apparent. In the two-layered defense signaling mechanisms, extracellular pattern-recognition receptors (PRRs) recognize conserved pathogen-associated molecular patterns (PAMPs) such as flagellin, chitin, or lipopolysaccharides and form the first line of basal defenses, also known as PAMP-triggered immunity (PTI) (Jones and Dangl, 2006; Schwessinger and Ronald, 2012). Elicitation of PTI leads to rapid increase in intracellular Ca^2+^, production of reactive oxygen species (ROS) and requires the defensive hormone salicylic acid (SA) to transduce the activation of downstream defense-associated genes such as *FLG22-INDUCED RECEPTOR LIKE KINASE 1* (*FRK1*), *WALL-ASSOCIATED KINASE 2* (*WAK2*), and *PATHOGENESIS-ASSOCIATED PROTEINS* (*PR1, PR2*) (Seybold *et al*., 2014; Bigeard *et al*., 2015; Sardar *et al*., 2017). Treatment of Arabidopsis suspension cells with SA decreases cellular pools of PtdIns with parallel increase in PtdIns4K and PtdIns(4,5)P_2_ levels (Krinke *et al*., 2007). Likewise, elicitation of PTI during *PstDC3000* exposure causes elevated InsP_3_ in Arabidopsis seedlings (Hung *et al*., 2014). Murphy *et al* (2008) reported the essentiality of InsP_6_ in maintaining basal defence against various pathogens. In their assays, *ipk1-1* and *mips2*, but not *mips1*, plants were compromised in PTI to *PstDC3000* (Murphy *et al*., 2008). Defenses against the cyst nematode is also reduced in the *ipk1-1* plants (Jain, 2015). Recently, it was shown that while *ipk1-1* plants remain deficient in induction of local resistance, PTI hallmarks such as intracellular Ca^2+^ spike, ROS production, and upregulated expression of responsive markers remained comparable to wild-type plants (Poon *et al*., 2020). An earlier study though contrastingly claimed that induction of PTI markers upon PAMP-treatment are deficient in *ipk1-1* (Ma *et al*., 2017). In the second immune layer, plants deploy intracellular nucleotide-binding leucine-rich repeat (NB-LRR) also known as resistance (R) proteins to sense activities of secreted effectors that attempt to thwart PTI. These recognitions lead to effector-triggered immunity (ETI) consisting mostly of PTI patterns but with stronger amplifications (Thomma *et al*., 2011; Cui *et al*., 2015). Effectors that trigger ETI are known as avirulent. AvrRpm1 or AvrRpt2, two *Pseudomonas* avirulent effectors cause PLC-dependent accumulation of phosphatidic acid (PA) in the presence of the cognate R genes *RPM1* or *RPS2*, respectively. As a further processed product of DAG, PA potentiates immune signaling (Andersson *et al*., 2006). While from the above studies it is increasingly obvious that InsP derivatives are vital for immune processes, their contributions or individual specificities are not known.

Here, we utilized several Arabidopsis InsP-biosynthesis/metabolism mutants to test their defensive potentials against the virulent *PstDC3000* strain. Primarily, among the InsP-kinase mutants investigated, we identify the role of IPK1, ITPK1 and VIH2 as suppressors of SA-dependent immunity. Constitutive activation of SA-dependent defenses confers enhanced basal immunity to *ipk1-1, itpk1-2* or *vih2-4* plants, and is abolished when the SA-signaling networks are disrupted. With comparative InsP profiling we reveal possible link between low InsP_8_, in these mutants to their elevated defenses. Together with the recently implicated roles of the above InsP-kinases in phosphate homeostasis, we demonstrate its intersection on SA-based immunity. Overall, through genetic and molecular evidences we reveal unique coordination between phosphate and defense signaling pathways highlighting responsive balances of a plant during stress adaptations.

## RESULTS

### IPK1, ITPK1 and VIH2 are suppressors of basal immunity against *PstDC3000*

Previously, it was reported that InsP_6_ is essential for promoting PTI and ETI in Arabidopsis (Murphy *et al*., 2008). The *ipk1-1* plants displayed deficient defenses against virulent and avirulent *PstDC3000* strains although neither basal SA nor its upregulation upon PAMP-treatment was affected thus suggesting immune compromises in processes downstream of SA (Murphy *et al*., 2008; Poon *et al*., 2020). Because *ipk1-1* plants are also depleted for InsP_7_ and InsP_8,_ whereas InsP_4_ accumulates (Stevenson-Paulik *et al*., 2005; Laha *et al*., 2015), their contributions to altered defense outcome remained uninvestigated. In order to gain deeper insights into this, we screened a series of InsP mutants for basal defenses against virulent *PstDC3000*. We chose T-DNA insertional mutants of *myo-INOSITOL KINASE* (*MIK*) (*mik-1*)(Kim and Tai, 2011), *INOSITOL 1,4,5-TRISPHOSPHATE KINASES* (*IPK2α*and *IPK2β)* (*ipk2α-1* and *ipk2β-1*) (Stevenson-Paulik *et al*., 2005), *INOSITOL 1,3,4-TRISPHOSPHATE 5/6-KINASES* (*ITPK1,3,4*) (*itpk1-2, itpk3-1, itpk4-1*) (Wilson and Majerus, 1997; Sweetman *et al*., 2007; Kim and Tai, 2011; Laha *et al*., 2020), and *DIPHOSPHOINOSITOL PENTAKISPHOSPHATE KINASES* (*VIH1,2*) (*vih1-1, vih2-4*) (Laha *et al*., 2015). The *ipk1-1* plants were used as a comparative control. To note, among these mutants, *mik-1, ipk1-1, ipk2β-1, itpk1-2* and *itpk4-1* are designated *lpa* mutants (Stevenson-Paulik *et al*., 2005; Kim and Tai, 2011). Seeds were obtained from Arabidopsis stock centre or as a gift and confirmed via genotyping PCR to identify homozygous insertions for the corresponding T-DNAs. In our growth regime, *ipk1-1* and *itpk1-2* plants displayed contrasting growth retardation phenotype with leaf epinasty (Figure 1a) as previously described (Stevenson-Paulik *et al*., 2005; Kuo *et al*., 2014; Kuo *et al*., 2018). These phenotypic defects were more pronounced under long-day (LD) conditions of growth. In the respective complemented lines *ipk1-1:Myc-IPK1* (Walia *et al*., 2020) or *itpk1-2:ITPK1-GFP* (Laha *et al*., 2020), growth defects were abolished and the transgenic plants were indistinguishable to Col-0 (Figure 1a; Figure S1). Further, constitutive expression of phosphate-starvation-inducible (PSI) genes and resulting elevated Pi accumulations previously known for *ipk1-1* or *itpk1-2* (Kuo *et al*., 2014; Kuo *et al*., 2018) were also restored to Col-0 levels in the complemented lines (Figure S2a) (Laha *et al*., 2020; Walia *et al*., 2020). Overall, these results validated the corresponding T-DNA mutational consequences on the abnormal growth phenotypes and constitutive PSR in *ipk1-1* or *itpk1-2*.

**FIGURE 1.**
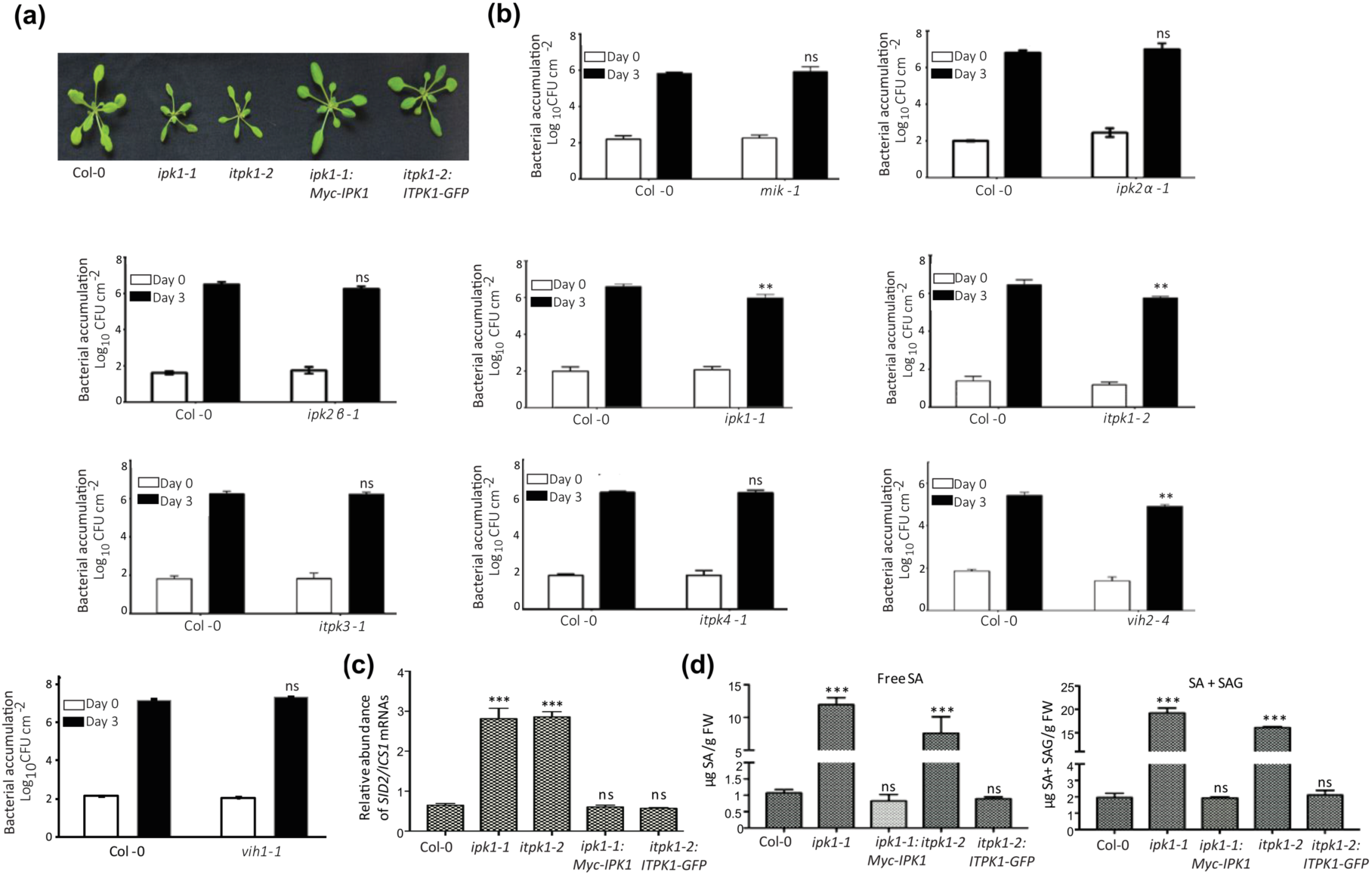
Selective Arabidopsis InsP-kinase mutants display enhanced basal defenses against *PstDC3000*. (a) Growth phenotypes of 3 week-old Col-0, *ipk1-1, itpk1-2*, and the corresponding complemented lines *ipk1-1*:*Myc-IPK1* and *itpk1-2*:*ITPK1-GFP*. (b) *PstDC3000* infection assays. Leaves from indicated mutants and Col-0 plants were infiltrated with bacterial suspension at a density of 5 × 10^4^ cfu/ml. Bacterial inoculum in the infiltrated leaves (Day 0) and its growth (Day 3) were quantified on media plates containing appropriate antibiotics. (c) Relative expression of *SID2*/*ICS1* in the indicated plants. Total RNA was isolated from leaves of 3-4 plants, reverse transcribed, and qRT-PCRs performed. Expressions were normalized to *MON1*. (d) Endogenous levels of free SA and total SA+SAG in the indicated plants. SA estimations were performed on 3-4 week-old plants. Data presented are mean (± SD) of three biological and technical replicates. Statistical analysis is according to Student’s *t*-test (\*\**p*<0.001, \*\*\**p*<0.0001, ns = not significant).

When challenged with the virulent *PstDC3000*, we noted a clear decrease in bacterial accumulations in *ipk1-1, itpk1-2* and *vih2-4*, but not other InsP-kinase mutants, amounting to reductions by ∼5-10 fold compared to Col-0 (Figure 1b). Whereas elevated basal immunity in *itpk1-2* and *vih2-4* were not known before, enhanced resistance of *ipk1-1* we observed in our assays in particular contradicts its increased susceptibility reported earlier (Murphy *et al*., 2008; Poon *et al*., 2020).

### Basal elevations in SA-based defenses associate with *ipk1-1, itpk1-2* and *vih2-4* plants

The above contrasting results led us to further investigate whether defense-associated features linked to enhanced basal immunity are represented in the above InsP-kinase mutants. As SA is key mediator of these defenses, we measured the relative transcript levels of *SALICYLIC ACID-DEFICIENT 2/ ISOCHORISMATE SYNTHASE 1* (*SID2*/*ICS1*), the enzyme performing defense-induced SA biosynthesis (Nawrath and Métraux, 1999). In *ipk1-1* or *itpk1-2*, but not their respective complemented lines, and also in the *vih2-4* plants strong upregulation in *SID2*/*ICS1* expressions relative to Col-0 were noted (Figure 1c; Figure S3a). Further, free or glucosyl moiety-conjugated SA (SAG) levels were higher in *ipk1-1, itpk1-2* or *vih2-4* extracts relative to Col-0 (Figure 1d; Figure S3e). These observations were at discrepancy with the earlier report (Murphy *et al*., 2008) showing that basal SA levels or its induction post-avirulent *PstDC3000* infection remains comparable between Col-0 and *ipk1-1*. Regardless, SA or SAG enhancements were restored to Col-0 levels in the respective complemented lines of *ipk1-1* or *itpk1-2* affirming that the SA elevations are indeed attributed to the corresponding *ipk1-1* or *itpk1-2* mutations.

Elevated SA induces *PR1* or *PR2* expressions. In *ipk1-1, itpk1-2* or *vih2-4* plants, prominent increase in *PR1*/*PR2* expressions with parallel accumulation of respective proteins in comparison to Col-0 were observed (Figure 2a,b; Figure S3). Endogenous levels of several PAMP-responsive genes such as *FRK1, WAK2, FOX, CYP81F2*, or *WRKY38* (Sardar *et al*., 2017) were also elevated in these mutants relative to Col-0 (Figure S3; Figure S4a,b). NON-EXPRESSOR of PATHOGENESIS-RELATED 1 (NPR1) orchestrates the induction of SA-responsive genes including *PR1* (Cao *et al*., 1997; Zhang *et al*., 1999). We detected clear enhancement in NPR1 protein levels in *ipk1-1* or *itpk1-2* than Col-0 (Figure 2c). This was in accordance to increased NPR1 accumulation upon SA-analog 2,6-dichloroisonicotinic acid (INA) treatments reported earlier for Col-0 (Kinkema *et al*., 2000). Basal defenses require ENHANCED DISEASE SUSCEPTIBILITY 1 (EDS1) and an *eds1* mutant (*eds1-2*) is hypersusceptible to virulent *PstDC3000* (Feys *et al*., 2001). Notably, in several auto-immune mutants EDS1 or the R protein SUPPRESSOR OF npr1-1 CONSTITUTIVE 1 (SNC1) accumulations are increased due to elevated SA responses (Garcia *et al*., 2010; Li *et al*., 2010; Dnyaneshwar Ingole *et al*., 2020). Distinct increase in EDS1 and SNC1 protein levels were observed in *ipk1-1* or *itpk1-2* compared to Col-0 (Figure 2d,e). As again, extracts from complemented lines restored EDS1 elevations to Col-0 levels. Further, increased callose deposits also signify induced basal defenses in auto-immune plants (Dnyaneshwar Ingole *et al*., 2020; Voigt, 2014). In *ipk1-1* or *itpk1-2*, endogenous callose accumulations were significantly higher than Col-0 providing further validation to their increased basal immunity we observed (Figure 2f). Taken together, our results convincingly illustrate auto-immune phenotypes of *ipk1-1, itpk1-2* and *vih2-4*.

**FIGURE 2.**
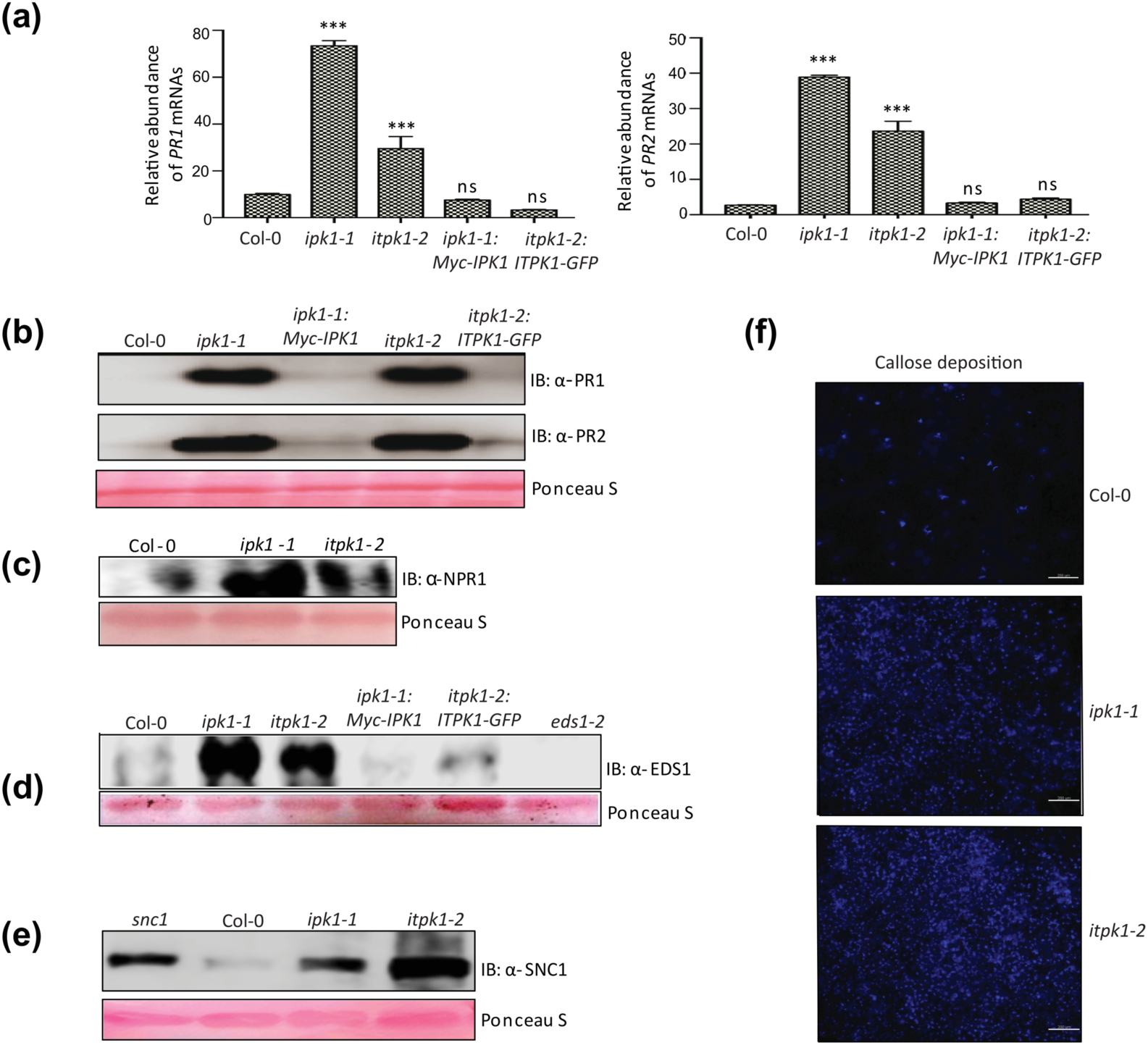
Expression of defense-associated genes/proteins are upregulated in *ipk1-1* or *itpk1-2* plants. (a) Relative expression of *PR1* or *PR2* transcripts. qRT-PCRs were performed and normalized to *MON1* gene-expressions as an internal control. Values are mean ± SD with 3 biological and technical replicates (n=3). Student’s *t*-test analysis is shown (\*\*\**p*<0.0001, ns = not significant). Immunoblots of defense-associated proteins (b) PR1 or PR2, (c) NPR1, (d) EDS1, or (e) R protein SNC1 in indicated plants. Total proteins were probed with respective antibodies. The membrane was stained with Ponceau S to demonstrate comparable protein loadings. (f) Callose deposits on Col-0, *ipk1-1*, or *itpk1-2* leaves. Fully expanded leaves from 3-week-old plants were stained for callose and observed under a fluorescence microscope. Scale bar = 50μm.

### SA-signaling routes are responsible for enhanced basal immunity in *ipk1-1* or *itpk1-2*

To genetically associate enhanced SA-regulated networks to heightened basal defenses in *ipk1-1* or *itpk1-2*, we generated *ipk1-1 eds1-2, ipk1-1 sid2-1, itpk1-2 eds1-2* and *itpk1-2 sid2-1* double mutants. An *eds1-2* plant has truncation in the *EDS1* gene rendering it non-functional whereas the *sid2-1* harbors a null-mutation in *SID2*/*ICS1* (Wildermuth *et al*., 2001, Cui *et al*., 2017). Loss of either *EDS1* or *SID2*/*ICS1* did not affect the growth retardation phenotypes of *ipk1-1* or *itpk1-2* implying that these defects are SA-independent (Figure 3a). Nonetheless, *eds1-2* and more prominently *sid2-1* mutation in *ipk1-1* or *itpk1-2* reduced the elevated *PR1* expressions (Figure 3b). Taking into account that EDS1 mediates positive amplification loop of SA-based defense signaling, residual *PR1* levels that remain elevated in *ipk1-1 eds1-2* or *itpk1-2 eds1-2* are likely indicative of this. Likewise, *FRK1* expression enhancements in *ipk1-1* was abolished by *sid2-1* but only modestly affected by *eds1-2* (Figure S4c,d). Upon challenge with *PstDC3000, ipk1-1 eds1-2, ipk1-1 sid2-1, itpk1-2 eds1-2* or *itpk1-2 sid2-1* plants lost enhanced basal immunity and instead were as hypersusceptible as *eds1-2* or *sid2-1* plants (Figure 3c). The results thus implied that primed defenses in *ipk1-1* and *itpk1-2* were indeed routed through SA-signaling networks. For additional support, we also generated *ipk1-1 npr1-1* double mutants. As observed earlier for *ipk1-1 eds1-2* or *ipk1-2 sid2-1* plants, growth defects of *ipk1-1* remained unchanged in *ipk1-1 npr1-1* reemphasizing that processes that are SA-independent likely cause the developmental deficiencies (Figure S5a). Loss of NPR1 resulted in substantial down-regulation of *PR1*/*PR2* transcript elevations in *ipk1-1* (Figure S5b). Overall, these evidences provide strong indications of *IPK1* and *ITPK1* functions as suppressors of SA-mediated immunity.

**FIGURE 3.**
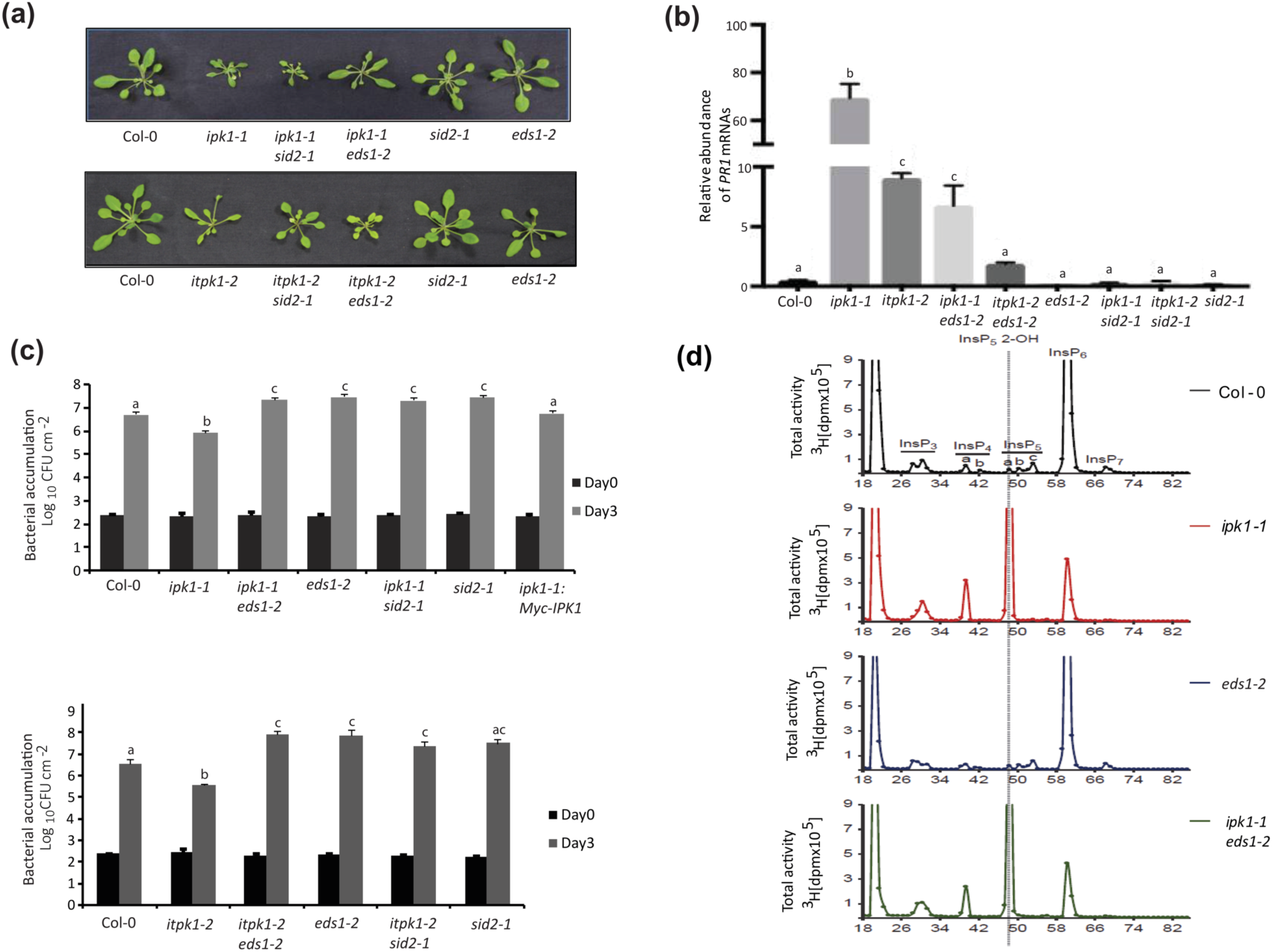
Loss of *SID2*/*ICS1* or *EDS1* abolishes enhanced basal immunity but not growth deficiencies of *ipk1-1* or *itpk1-2* plants. (a) Growth phenotypes of 3 week-old Col-0, *ipk1-1, itpk1-2, ipk1-1 eds1-2, ipk1-1 sid2-1, itpk1-2 eds1-2, itpk1-2 sid2-1, sid2-1*, and *eds1-2* plants. (b) Relative expression of *PR1* in the indicated plants. qRT-PCR was performed on cDNAs generated from above plants. As an internal control, *MON1* gene expression was used. Expressions are presented as mean ± SD (n=3). (c) *PstDC3000* growth assays on *eds1-2* or *sid2-1* combinatorial mutants with *ipk1-1* or *itpk1-2*. Pathogenesis assays were performed and quantified as earlier. Statistical significance shown with different letters is according to posthoc Tukey’s test (p<0.05). (d) Incorporating *eds1-2* does not change the InsP profiles of *ipk1-1*. SAX-HPLC profiles of Tritium-labeled InsPs in Col-0, *ipk1-1, eds1-2*, and *ipk1-1 eds1-2* extracts are shown. Labeling was performed on 3-week-old plants and activities determined by scintillation counts of fractions containing InsP_2_-InsP_8_.

### Reduced InsP_8_ may contribute to elevated defences of *ipk1-1* and *itpk1-2*

Primarily, IPK1 and ITPK1 activities constitute a metabolic pair that coordinates synthesis of InsP_7_ from InsP_5_ (Laha *et al*., 2019; Adepoju *et al*., 2019; Whitfield *et al*., 2020). InsP_7_ is further pyrophosphorylated by VIH1/2 to produce InsP_8_ (Laha *et al*., 2015). Along with reduced InsP_7_ in *ipk1-1* or *itpk1-2*, InsP_8_ levels are also lower in these mutants similar to *vih2-4* (Stevenson-Paulik *et al*., 2005; Laha *et al*., 2015; Kuo *et al*., 2018). The *vih2-4* plants however have basal levels of InsP_6_ but elevated InsP_7_ (Laha *et al*., 2015). In *itpk4-1*, low InsP_6_ have been recently reported but levels of InsP_7_ in these plants remain contrasting between two published studies (Kuo *et al*., 2018; Laha *et al*., 2020). Total InsP profiles of *mik-1* or *ipk2β-1* especially in context of InsP_7_ or InsP_8_ is not known. In order to determine which InsPs contribute towards enhanced SA-dependent immunity, we performed SAX-HPLC analyses of [^3^H]-inositol-labelled seedlings of Col-0, *mik-1*, and *ipk2β-1*. We also included *ipk1-1, itpk1-2* and *itpk4-1* in these measurements. Although InsP_6_ levels were reduced in *mik-1* or *ipk2β-1* seedlings, InsP_7_ or InsP_8_ levels were unchanged from Col-0 (Figure 3d and Figure S6). From these analysis, we surmised a possible correlation between specific InsP and altered defenses. Reduced InsP_6_ is not likely the cause of enhanced basal immunity since *mik-1, ipk2β-1* or *itpk4-1* display Col-0-comparable defenses. Neither the decreased InsP_7_ is a cause for the same since *itpk4-1* with claimed deficiency in this InsP (Kuo *et al*., 2018) does not show altered defensive capabilities against *PstDC3000*. Increased InsP_7_ levels in *vih2-4* also does not result in enhanced susceptibility, as would be expected from this hypothesis. Taken together, by exclusion it is suggestive that low InsP_8_ common to *ipk1-1, itpk1-2* and *vih2-4* may be linked to their enhanced basal immunity. Interestingly, InsP separation profile of *ipk1-1eds1-2* mirrored *ipk1-1* patterns indicating that increased SA-based defenses were downstream consequences to the altered InsPs (Figure 3d).

### Constitutive PSR augments enhanced SA-defenses in *ipk1-1* and *itpk1-2*

Constitutive activation of PSI-genes is characteristic of *ipk1-1* or *itpk1-2* (Kuo *et al*., 2014; Kuo *et al*., 2018). Under sufficient phosphate conditions, the transcription factor PHOSPHATE STARVATION RESPONSE 1 (PHR1) is sequestered by SPX1 (SYG1/Pho81/XPR1-domain containing protein 1) in the presence of InsP_8_ cofactor (Puga *et al*., 2014; Zhong *et al*., 2018). This prevents its binding to promoters of PSI-genes and activating PSR. When soil Pi is low, phosphatase activity of VIH2 is activated resulting in reduced InsP_8_, liberating PHR1 and causing 26S proteasome mediated degradation of SPX1 (Zhu *et al*., 2019, Ried *et al*., 2019). In a breakthrough report, PHR1 was shown to suppress several SA-responsive gene expressions (Castrillo *et al*., 2017). Arabidopsis *phr1 phl1* double mutant (*PHL1* is a paralogue of *PHR1*) is enhanced resistant to *PstDC3000*. While reduced InsP_8_ likely activates PHR1/PHL1 functions in *ipk1-1* or *itpk1-2*, increased SA-dependent immunity in these plants appear as a contradiction. To determine whether constitutive PSR influences enhanced SA-responses of *ipk1-1* or *itpk1-2*, we obtained *ipk1-1 phr1 phl1* (Kuo *et al*., 2018) and generated *itpk1-2 phr1 phl1* triple mutants by genetic crossing of *phr1 phl1* double mutant with *itpk1-2*. A *phr1 phl1* double mutant is defective in PSR induction and basal levels of Pi are lower than Col-0 (Bustos *et al*., 2010; Kuo *et al*., 2014). We observed that growth defects of *ipk1-1* or *itpk1-2* are not recovered in *ipk1-1 phr1 phl1* or *itpk1-2 phr1 phl1*, respectively suggesting that these phenotypes are independent of PHR1/PHL1 (Figure 4a). Further, as expected increased endogenous Pi levels in *ipk1-1* or *itpk1-2* and elevated expression of PSI-genes *IPS1* or *SPX1* are reduced below Col-0 levels in the above triple mutants (Figure S2a,b) (Kuo *et al*., 2014).

**FIGURE 4.**
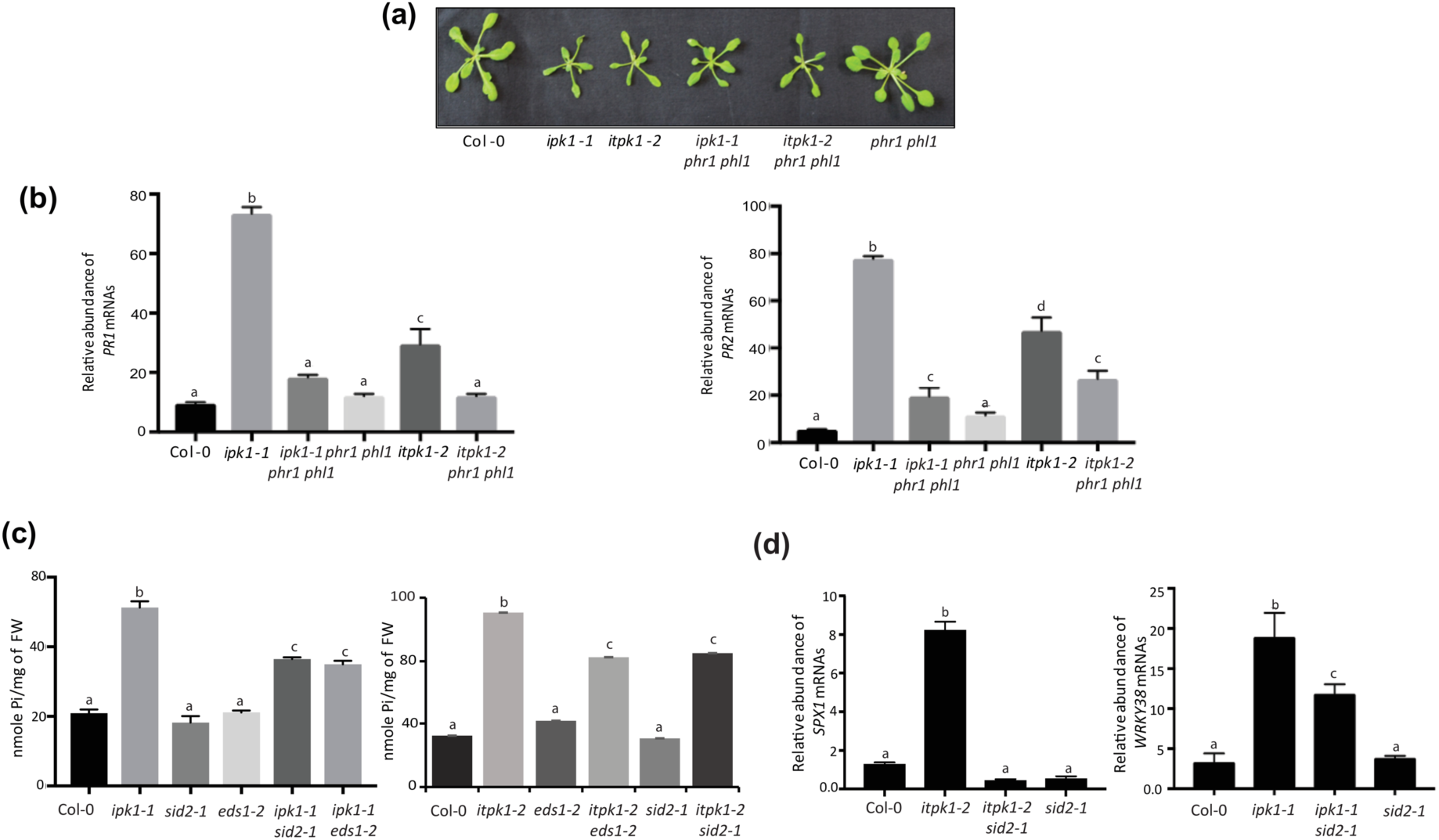
Enhanced basal defenses but not phenotypic growth defects in *ipk1-1* or *itpk1-2* are partially *PHR1*/*PHL1*-dependent. (a) Growth phenotypes of 3 week-old Col-0, *ipk1-1, itpk1-2, ipk1-1 phr1 phl1, itpk1-2 phr1 phl1*, and *phr1 phl1* plants. (b) Introducing *phr1 phl1* mutations dampen elevated *PR1*/*PR2* expressions in *ipk1-1* or *itpk1-2*. (c) Elevated endogenous Pi, or (d) heightened *SPX1* or *WRKY38* expression in *ipk1-1* or *itpk1-2* are downregulated by *eds1-2* or *sid2-1* mutation. qRT-PCR are normalized to expression of the internal control *MON1* gene. All data presented here are mean values ± SD of three biological and technical replicates (n=3). Different letters indicate differences according to posthoc Tukey’s test (p<0.05).

To determine whether abolishment of enhanced PSR by the loss of *PHR1*/*PHL1* affected basal accumulation of *PR1*/*PR2* transcripts, we compared their relative expression levels between the respective triple and the single InsP-kinase mutants. In accordance with the enhanced defenses against *PstDC3000* infection reported earlier (Castrillo *et al*., 2017), both *PR1* and *PR2* expressions were slightly but not significantly elevated in *phr1 phl1* plants when compared to Col-0 (Figure 4b). Surprisingly, introducing *phr1 phl1* mutation in *ipk1-1* or *itpk1-2* substantially reduced the upregulated *PR1*/*PR2* expressions (Figure 4b). These results suggested that elevated PSR associated with PHR1/PHL1 activities contributes positively to the enhanced defense-gene expressions in the above InsP-kinase mutants.

### Elevated SA in *ipk1-1* or *itpk1-2* aggravate PSR via feedback mode

Several PSI-genes have SA-inducible elements in their promoters (Baek *et al*., 2017). Therefore, to determine whether elevated SA reciprocates on PSR in *ipk1-1* and *itpk1-2*, we investigated for changes in endogenous Pi elevations in *ipk1-1 sid2-1* and *itpk1-2 sid2-1* plants. Loss of *SID2*/*ICS1* or *EDS1* toned-down elevated Pi levels in *ipk1-1* or *itpk1-2* (Figure 4c). When compared for relative expression of PSI-genes, *SPX1* or the transcription factor *WRKY38* displayed reduced expressions in *itpk1-2 sid2-1* or *ipk1-1 sid2-1*, respectively compared to the *itpk1-2* or *ipk1-1* parent (Figure 4d). Likewise, *ipk1-1 npr1-1* plants also contained lower endogenous Pi-levels than *ipk1-1*, and displayed complete or partial restoration of *WRKY38* or an another PSR-marker *SAL2* (3’(2’), 5’-bisphosphate nucleotidase/inositol polyphosphate 1-phosphatase) (Gil-Mascarell *et al*., 1999) expressions, respectively to Col-0 levels (Figure S5d). Interestingly, it is known that *WRKY38* is SA-inducible and requires NPR1 for transcriptional activation (Kim *et al*., 2008). With these results, it is suggestive that heightened SA-signaling sectors aggravate but are not the direct cause of PSR in *ipk1-1* or *itpk1-2*.

To further substantiate the SA-promotion on PSI-gene expression, we treated Col-0 with SA and measured the kinetics of *SPX1* or *WRKY38* transcripts. Remarkably, SA exposure progressively increased *SPX1* or *WRKY38* expressions from 30-120 min post-treatment (mpt) (Figure 5a). The kinetics closely matched those of *PR1* inductions in the same samples. These data therefore corroborated that indeed SA can potentiate PSR in plants by directly activating the transcription of at least a subset of PSI-genes.

**FIGURE 5.**
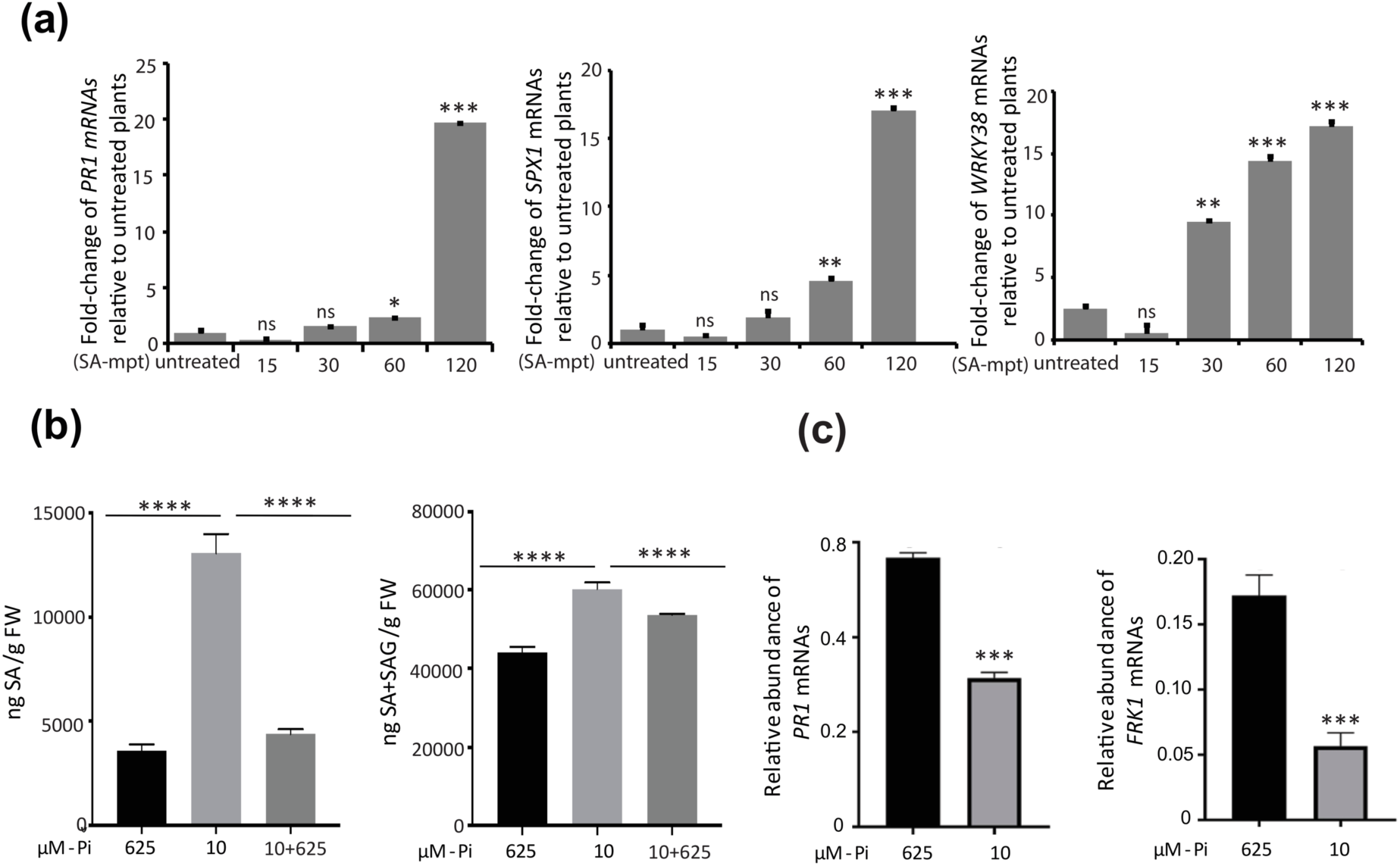
SA application induces PSI-gene expressions. SA elevations associate with PSR although defense-gene expressions are suppressed. Kinetics of (a) *PR1, SPX1*, or *WRKY38* expressions post-SA-treatment on Col-0. Approximately, 3-4 week-old Col-0 plants were spray-treated with SA and samples harvested at indicated time points (mpt, minutes post-treatment) and processed for qRT-PCR analysis. (b) Free SA and total SA+SAG levels in Col-0 under Pi-sufficient (625 μM-Pi), Pi-deprived (10 μM-Pi), or Pi-replenished post-starvation (10-625 μM-Pi) conditions. (c) Relative expression levels of *PR1* or *FRK1* genes under Pi-sufficient or Pi-deprived growth. For these assays, Col-0 seedlings grown for 14-days under Pi-sufficient conditions were subjected for 4-days to Pi-starvation and then re-supplemented with replete Pi levels for 12 hrs, before being processed. The qRT-PCR data are normalized *MON1* gene expressions. All data shown here are mean values (± SD, n=3) with Student’s *t*-test for statistical analysis (**p<0.001, ***p<0.0001,****p<0.00001).

### PSR reprogrammes defense-signaling networks of SA

In *phr1 phl1* plants, a significant proportion of differentially expressed genes are SA-inducible (Castrillo *et al*., 2017). These include immune-associated genes that possess P1BS (PHR1-binding site) elements and therefore are direct PHR1-targets. If indeed SA is augmenting PSR as we noted in *ipk1-1* or *itpk1-2* plants, it is possible that its defensive roles are re-routed to support PSR processes. To investigate this, we subjected Col-0 plants to Pi-starvation and measured the basal levels of SA and expression of downstream *PR1, FRK1*, or *NPR1* genes. Under Pi-deprivation, SA or SAG levels were significantly elevated and restored rapidly when exogenously supplemented with phosphate in the growth medium (Figure 5b). Most surprisingly, even with SA-elevations, expression of *PR1* or *FRK1* was downregulated upon Pi-starvation supporting suppression of basal immunity as proposed earlier (Figure 5c) (Castrillo *et al*., 2017). *WRKY38* is a known negative regulator of basal immunity (Kim *et al*., 2008). Endogenous *PR1* expression levels are higher in *wrky38-1* plants and result in enhanced immunity to *PstDC3000*. Thus, with these into consideration it is implicative that although PSR induces SA, its roles are diverted from immune promotion to PSR supportive functions possibly involving WRKY38.

### PHR1-dependent induction of PSI-genes affect intensity of PTI responses

Implications on positive influence of SA on expression of PSI-genes, prompted us to determine whether in a physiological context of a pathogen threat a similar phenomenon is observed. Towards this, Col-0 or *phr1* plants were challenged with *PstDC3000* as earlier and hallmarks of PSR responses tested in the infected tissues. As is expected, expression of *SID2*/*ICS1* and *PR1* was induced in Col-0 at 3-dpi (Figure 6a,b). Curiously, these upregulations in the *phr1* plants were significantly lower than Col-0. Whereas *SID2*/*ICS1* or *PR1* transcripts showed >10-fold increase in Col-0, these were only ∼6-fold elevated in *phr1* plants post-infection. Further, while expression of PSI-genes *PHT3;2* or *PAP17* demonstrated clear upregulations with *PstDC3000* exposure on Col-0, their induction in *phr1* plants were comparatively lower (Figure 6c,d). Likely, functional PHL1 in *phr1* plants accounted for the residual elevation. Not the least, we surprisingly noted clear increase in endogenous Pi levels in Col-0 upon *PstDC3000* infection suggesting that PTI elicitation is accompanied by changes in Pi-homeostasis possibly through induced PSR pathway (Figure 6e). Indeed, in *phr1* plants this increase was barely detected suggesting that these responses are primarily PHR1-orchestrated. We recently reported that IPK1 and ITPK1 protein stabilities are increased during PSR perhaps signifying their intersection on diverse signaling pathways (Walia *et al*., 2020). Likewise, upon *PstDC3000* challenge on *ipk1-1*:*Myc-IPK1* or *itpk1-2*:*ITPK1-GFP* plants we noted improvements in IPK1 and ITPK1 protein stabilities likely reflective of a similar phenomenon (Figure 6f). With these results, we reveal regulated synergism between basal defenses and phosphate homeostasis incorporating roles of selective InsP-kinases, IPK1 and ITPK1. Overall, with our investigations here we identify intricacies of signaling crosstalks, modulations, and re-adjustments during stress adaptations.

**FIGURE 6.**
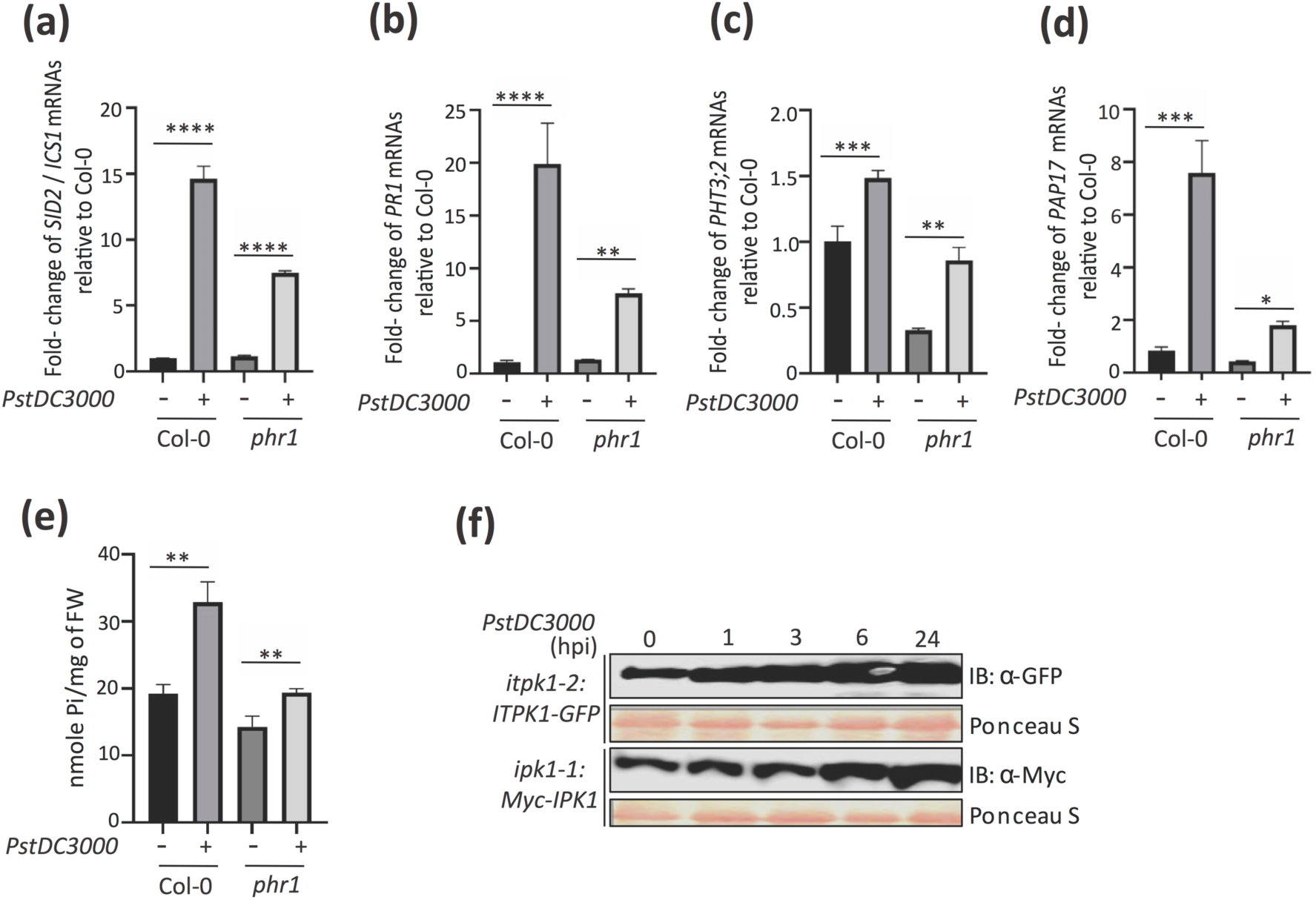
*PHR1* modulates induction of defense gene expressions during basal immunity. Relative expression levels of (a) *SID2*/*ICS1*, (b) *PR1*, (c) *PHT3;2*, (d) *PAP17* in Col-0 and *phr1* plants challenged with *PstDC3000* at 3dpi (days post-infection). Expression levels are normalized to *MON1* and reported as fold change relative to Col-0 (e) Changes in endogenous Pi levels in Col-0 and *phr1* plants at 3dpi. Data shown here are mean values (± SD, n=3). Statistical significance is with Student’s *t*-test (\*\**p*<0.001,\*\*\**p*<0.0001,**** *p*<0.00001). (f) Kinetics of IPK1 and ITPK1 protein levels after *PstDC3000* infections at indicated time-points (hours post-infection, hpi). Immunoblots were performed with anti-GFP (for ITPK1-GFP) or anti-Myc (for Myc-IPK1) antibodies. PonceauS staining of the membranes denote comparable protein loadings.

## DISCUSSION

Plant immune responses recruit complex signaling networks and intricate interplay of phytohormone activities. While these remains tightly orchestrated and transitory under normal physiological threats to avoid fitness costs, the same is not always apparent in several characterized autoimmune mutants. With the pleiotropic phenotypes displayed by these mutants the complexity of crosstalks that occur among different signaling pathways and the players/signals that mediate these are increasingly being unraveled (van Wersch *et al*., 2016). Our study here especially highlights the selective involvement of two InsP-kinases IPK1 and ITPK1 in SA-dependent regulation of innate immune signaling. With previously established roles in maintaining Pi-homeostasis (Kuo *et al*., 2014; Kuo *et al*., 2018), JA (Sheard *et al*., 2010; Mosblech *et al*., 2011; Boss & Im, 2012), and auxin signaling (Tan *et al*., 2007; Laha *et al*., 2020) we demonstrate their repercussions on SA-mediated basal defenses. Both *ipk1-1* and *itpk1-2* plants, similar to *vih2-4*, display enhanced immunity to virulent *PstDC3000*. The common InsP_7_/InsP_8_ reduction in these mutants posits that perturbations leading to their enhanced immunity may be attributed to multiple factors. Suppression of SA-driven pathway by the loss of either *SID2*/*ICS1* or *EDS1* mitigates the enhanced resistance in *ipk1-1* or *itpk1-2* plants implying that the immune disturbances are likely upstream of SA-biosynthesis as evidenced by increased expression of *SID2*/*ICSI* in these mutants relative to Col-0.

It is evident that our results present contradictions to earlier reports claiming the essentiality of InsP_6_ in maintaining basal defenses against *PstDC3000* (Murphy *et al*., 2008; Ma *et al*., 2017; Poon *et al*., 2020). First, with the extensive list of InsP-kinase mutants we tested in our pathogenesis assays, we are unable to correlate reduced InsP_6_ levels to altered basal immunity. Especially *mik-1, ipk2β-1* or *itpk4-1* have lower cellular InsP_6_ pools but display Col-0-comparable PTI. While we confirm that InsP_6_ is reduced in *ipk1-1* as earlier (Murphy *et al*., 2008; Kuo *et al*., 2018), for *itpk1-2* we mirror Laha *et al* (2020) that phytic acid levels remains unchanged. Nevertheless, even after considering these InsP discrepancies, only *ipk1-1* and *itpk1-2* have elevated basal immunity in our assays. Therefore, as implied earlier perhaps InsP_6_ changes at specific cellular locale(s) and not globally *per se* may primarily affect basal immunity. We describe a possible scenario in the later section that may be in accordance with this hypothesis. Second, we detect prominent increase in endogenous SA levels causing elevated expression of *PR1*/*PR2*, several PTI markers, EDS1, SNC1 or NPR1 proteins in *ipk1-1* or *itpk1-2* plants. These upregulations are restored to Col-0 levels in the respective complemented lines thus providing support to our observations. Murphy *et al* (2008) however did not find noticeable change either in the basal SA levels or its increase upon avirulent pathogen challenge in *ipk1-1*. Further, although Poon *et al* (2020) did not measure the endogenous expression levels of *FRK1* or other PTI markers, their induction upon PAMP-treatment was not affected in *ipk1-1*. Ma *et al* (2017) in contrast showed impaired induction of *FRK1* and other PTI responses in *ipk1-1* (also in *mips1* and *mips2*) with PAMP-exposure. These increasing disparities between studies clearly raise considerable doubts of InsP_6_ role in basal immunity. Perhaps consorting factors such as growth regimen, soil compositions, light, temperature or humidity regulations impact the immunity in *ipk1-1* making the responses conditional.

The *vih2-4* plants have wild-type InsP_6_, although InsP_7_ or InsP_8_ levels are elevated or reduced, respectively (Laha *et al*., 2015). The hyper-defensive nature of *vih2-4* plants we observe may hold a possible clue to relate specific InsP changes to basal immune enhancements. That, *ipk1-1* or *itpk1-2*, but not other InsP mutants we tested here, share InsP_8_ deficiency to *vih2-4* (Laha *et al*., 2020) lends support our speculations. Antagonism between JA-SA crosstalks are well known and exploited intensely by both pathogens and plant hosts in their efforts to colonize or resist, respectively (Caarls *et al*., 2015). Coronatine, secreted by *P. syringae* pathovars is a JA-analog and attempts to suppress SA-mediated defenses by upregulating JA-responses (Zheng *et al*., 2012). NPR1 in turn is essential for inhibiting the expression of JA-responsive genes (Spoel *et al*., 2009). With InsP_8_ requirement as a cofactor *vih2-4* plants therefore are compromised in JA-mediated defenses and remain hypersusceptible to herbivory (Laha et al., 2015). Elevated expression of PTI-responsive genes/SA levels in *vih2-4* and likewise in *ipk1-1* or *itpk1-2* may hence be due to impaired JA signaling resulting in elevated SA-defenses. The observations from Poon *et al* (2020) that prior injection of air/water in *ipk1-1* enhances resistance to *PstDC3000* may be the result of JA-deficiencies causing SA-based immune elevations. Mosbelch *et al* (2011) data that *ipk1-1* plants are JA-hypersensitive and resist *Plutella xylostella* caterpillar feeding however argues against this interpretation. Increased basal immunity in *ipk1-1* or *itpk1-2* may alternatively be attributed to reduced InsP_7_ levels and through a different mechanism than JA-antagonism. As the recently refined cofactor for auxin signaling, InsP_7_ facilitates TRANSPORT INHIBITOR RESPONSE 1 (TIR1)-mediated degradation of AUX/IAA repressors (Laha *et al*., 2020). Considering exogenous application of auxin potentiates *PstDC3000* colonization by interfering with SA defenses (Navarro *et al*., 2006; Wang *et al*., 2007), auxin-signaling impairments may conversely enhance basal immunity. Indeed, the auxin-insensitive mutant *axr2-1* significantly reduces bacterial growth in SA-impaired *NahG* plants (Wang *et al*., 2007). Thus, a different mode of impairment in *ipk1-1* or *itpk1-2* than *vih2-4* may be the cause of similar consequence of upregulated SA-mediated defenses.

Under Pi-deplete conditions plant immunity is compromised for successful colonization of phosphate-remobilizing microbiota to enhance phosphate availability in soil for uptake (Castrillo *et al*., 2017). The upregulation of SA-inducible genes in PSR-deficient *phr1 phl1* plants suggests antagonism between PSR and SA responses. However, a more complex interplay we reveal here is likely deployed in the actual physiological responses to Pi-deprivation. We show that constitutive PSR is partly responsible for aggravating SA-dependent immunity in *ipk1-1* or *itpk1-2*. Loss of *PHR1*/*PHL1* from *ipk1-1* or *itpk1-2* in addition to alleviating the enhanced PSR/Pi levels also tones down *PR1/PR2* upregulations. Reciprocally, *sid2-1* or *eds1-2* mutations that abrogate SA-mediated defenses partially mitigate constitutive PSR in *ipk1-1* or *itpk1-2*. As predicted recently, several PSI-genes including *SPX1* harbor SA-responsive elements (Baek *et al*., 2017) and may utilize the SA elevations we observe during PSR for optimal expression. That exogenous SA induces *SPX1* and *WRKY38* expressions supports our claim. Interestingly, *WRKY38*, a negative regulator of basal immunity, is SA-inducible and requires NPR1 (Kim *et al*., 2008). Phosphite, salt-conjugates of phosphorous acid ([HPO_3_]^2-^) used routinely as a fungicide, primes SA-dependent immunity and suppresses expression of PSI-genes (Varadarajan *et al*., 2002; Achary *et al*., 2017).

Upregulated expression of several PSI-genes upon a pathogen challenge we demonstrate here possibly highlights the energy necessities of defense, a process to which PSR-promoted Pi-uptake may significantly contribute. Multiple studies indeed report significant increase in ATP/ADP ratio (also known as adenylate charge status) during increased phosphate uptake post Pi-starvation (Zhu *et al*., 2019; Riemer *et al*., 2020). Activities of IPK1-ITPK1 pair and VIHs especially are modulated by changes in adenylate charge in a cell (Zhu *et al*., 2019; Riemer *et al*., 2020; Whitfield *et al*., 2020). Taken together these results suggest a mutual modulatory loop between SA and PSR networks involving specific InsPs. In Pi-starved wild-type plants immune suppressions are possibly accomplished by blocking SA-transduction of immunity by involving *PHR1* functions as a negative regulator of defensive-associated genes (Castrillo *et al*., 2017). Induced SA are channeled via NPR1-dependent roles into processes such as *WRKY38* or *SPX1* expressions that promote PSR, simultaneously maintaining immune gene inhibitions. With possible dysfunctions in this coordination further aided by reduced InsP_8_ levels (or changes in other unidentified InsPs), *ipk1-1* or *itpk1-2* hence show constitutive PSR accompanied with elevated SA responses. The autoimmune *siz1-2* plants that exhibit constitutive SA and PSR responses may exemplify another cellular module that intersects on PSR-SA harmony (Miura *et al*., 2011).

With the above, it is encouraging to consider that a common protein hub regulated by IPK1/ITPK1 activities orchestrate the fine-tuning of cellular signaling logistics upon a stimulus. We recently in a remarkable parallel with earlier animal studies identified that IPK1-ITPK1 via strict locale-specific roles moderates the functioning of CONSTITUTIVE PHOTOMORPHOGENESIS 9 (COP9 also known as CSN) signalosome (Walia *et al*., 2020). We showed that CSN activities in turn maintain cellular Pi-homeostasis by regulating the Cullin RING Ligases (CRLs) functioning in targeted-degradation of substrates by the 26S proteasome complex. Functional deficiencies in CSN result in strong pleiotropic effects including auxin-insensitivity (Dohmann *et al*., 2008), constitutive activation of PSR (Walia et al., 2020), and impaired JA-defenses accompanied by elevated PR1 transcripts (Hind *et al*., 2011), thus revealing strong parallel with *ipk1-1* or *itpk1-2* defects. Rice SPX4, SNC1 or NPR1 stabilities are monitored by CRL functions placing molecular implications of IPK1/ITPK1 on defense outcomes via CSN-CRL dynamics (Gou *et al*., 2012; Ruan *et al*., 2019; Ried *et al*., 2019; Shen *et al*., 2020). These indications lead us to advocate that between the investigated InsP-kinases, IPK1 and ITPK1 have acquired unique localized roles that impinge on central cellular machineries impacting multiple signaling networks. In conclusion, with our studies here we decipher complexities of plant signaling that recruits seemingly antagonistic networks and reassigns roles to specific players to elicit stimulus-appropriate response outputs.

## Material and Methods

### Plant material and growth conditions

Most T-DNA insertional mutants of *Arabidopsis thaliana* (accession Col-0) for the investigated InsP-kinases were obtained from Arabidopsis Biological Resource Centre (ABRC; www.abrc.osu.edu). The details of *ipk1-1, itpk1-2, ipk2β-1, itpk4-1, vih1-1, vih2-4* have been described earlier (Stevenson-Paulik *et al*., 2005; Kim and Tai, 2011; Kuo *et al*., 2014; Laha *et al*., 2015; Kuo *et al*., 2018; Laha *et al*., 2020). Primers used in genotyping for homozygous T-DNA insertions are listed in Supplementary Table 1. Seeds were cold-stratified for 2-days in dark, surface sterilized with 30% bleach, washed with three rinses of sterile water and then germinated on 0.5x Murashige and Skoog (MS) plus 1% sucrose containing agar plates. Growth chambers were maintained at 24°C with 70% RH and long-day (LD) conditions (16hrs:8hrs; light: dark cycle) with light intensity 100 µmol µm^-2^s^-1^.

For Pi-starvation assays, plants grown on media plates as above were transferred 7-days post-germination to liquid MS medium in 12-well sterile culture plates containing either 625 µM (Pi-sufficient) or 10 µM (Pi-deficient) KH_2_PO_4_ for 4 days before being used for indicated assays.

### Generation of combinatorial mutants

To generate *ipk1-1 eds1-2, itpk1-2 eds1-2, ipk1-1 sid2-1, itpk1-2 sid2-1, ipk1-1 npr1-1* or *itpk1-2 phr1 phl1* combinatorial mutants, *ipk1-1* or *itpk1-2* were genetically crossed with *eds1-2* (Cui *et al*., 2017), *sid2-1* (Wildermuth *et al*., 2001), *npr1-1* (Ramírez *et al*., 2010), or *phr1 phl1* (Kuo *et al*., 2014) plants. Required combination of mutations and their homozygosity was determined by genomic PCRs of segregating plants either in the F2 or F3 populations. Primers for the same are listed in Supplementary Table 1.

### *In planta PstDC3000* growth assays

Bacterial growth assays for *PstDC3000* were performed as earlier (Bhattacharjee *et al*., 2011). In brief, leaves from indicated plants soil-grown for 3-4 weeks were infiltrated with *PstDC3000* strain at a bacterial density of 5×10^4^ cfu/ml using a needleless syringe. Bacterial accumulation was measured at 0 and 3 days post-infiltration (dpi) by harvesting leaf discs of defined diameter, macerating in 10 mM MgCl_2_, and plating serial dilutions on Pseudomonas agar plates containing with 25 µg/ml Rifampicin. Growth of bacteria in the infiltrated leaves are reported as Log_10_cfu/cm^2^. For qPCRs and Pi estimation, a bacterial inoculum of 10^6^ cfu/ml was used. Tissues were harvested before and after infiltration (3 dpi) and processed accordingly.

### Salicylic acid (SA) measurements or treatments

Free and glucose-conjugated SA were measured according to (DeFraia *et al*., 2008) employing the Acinetobacter sp. ADPWH_lux biosensor system. Extracts from *sid2-1* plants spiked with known amounts of SA was used to generate standard curve. Luminescence was measured in a POLARStar Omega Luminometer (BMG Labtech). Data presented here are mean (± SD) of three biological replicates.

To determine induction of PSI genes, Col-0 plants grown on soil for 3-4 weeks were spray-treated with 0.5 mM SA (Sigma-Aldrich) and tissues harvested at indicated time-points for further analysis.

### RNA extraction, cDNA synthesis and qRT-PCR analysis

Total RNA was extracted with RNAiso Plus (Takara) and cDNA synthesized with iScript cDNA Synthesis Kit (Bio-Rad) following manufacturer’s instructions. Primers used for qRT-PCR primers are listed in Supplementary Table 1. All qRT-PCRs were performed in QuantStudio 6 Flex Real-Time PCR system (Applied Biosystems) with 5X HOT FIREPol® EvaGreen® qPCR Mix Plus (ROX) (Solis BioDyne) according to the manufacturer’s instructions. Each experiment contained three biological and technical replicates (n=3). Arabidopsis *MON1* (*At2g28390*) expression was used as internal control (Bhattacharjee *et al*., 2011). Relative expression was calculated according to the PCR efficiency^-ΔΔCt formula. Expressions were normalized relative to levels in Col-0 and plotted as fold-change.

### Protein extraction and western blots

For immunoblotting, leaf tissues from indicated plants were homogenized in 6 M Urea, clarified by centrifugation at 16,000 rpm at 4°C, mixed with 2× Laemmli buffer (0.1M Tris pH 6.8, 20% w/v glycerol, 4% w/v SDS, 100 mM DTT and 0.001% w/v Bromophenol blue), separated on SDS-PAGE and transferred onto polyvinylidene fluoride (PVDF) membrane. Blocking was performed with 5% w/v non-fat skimmed milk powder in 1× TBST (Tris-Buffered Saline, 0.1% w/v Tween® 20 Detergent) for 1 hr, rinsed and incubated overnight at 4°C with indicated primary antibodies [anti-SNC1 (Abiocode; R3588-1), anti-PR1, anti-PR2, anti-NPR1 or anti-EDS1 (Agrisera; AS10687, AS122366, AS121854, AS132751, respectively)]. The membranes were washed the next day with three rinses of TBST, incubated at RT for 1 hr with secondary antibodies conjugated with horseradish peroxidase (HRP) (Santa Cruz Biotech). Rinsed membranes were then treated with ECL Prime solution (GE Healthcare) and luminescence imaged with an ImageQuant LAS 4000 system (GE Healthcare).

### Callose deposition assay

Callose staining was performed according to (Schenk and Schikora, 2015). Callose spots were observed under a Nikon fluorescence microscope equipped with a camera using a DAPI filter and UV light.

### Extraction of InsPs and SAX-HPLC profiling

Profiling of InsPs from indicated InsP-kinase mutants (and Col-0) were performed as according to (Laha *et al*., 2020). Briefly, Arabidopsis seedlings germinated for 10-days in 0.5x MS sucrose agar plates were transferred to liquid media containing 30 µCi mL-1 of [^3^H]-myo-inositol (30 to 80 Ci mmol-1; Biotrend) for 6 days. Seedlings were then rinsed with sterile water and flash-frozen with liquid N_2_. InsP extraction methodology was as described earlier (Azevedo and Saiardi, 2006). Extracts were then resolved by strong anion exchange high performance liquid chromatography (SAX-HPLC) using a Partisphere SAX 4.6 x 125 mm column (Whatman).

### Phosphate (Pi)-estimation assay

Total Pi measurements in the indicated plants were performed according to (Chiou *et al*., 2006). Plant tissues frozen with liquid N_2_ were homogenized in extraction buffer (10 mM Tris-HCl, 1 mM EDTA, 100 mM NaCl, 1 mM PMSF), mixed with 1% glacial acetic acid (ratio 1:9 v/v) and then incubated at 42°C for 30 mins. To the reaction, Pi-assay solution (0.35% NH_4_MoO_4_, 0.86 N H_2_SO_4_ and 1.4% ascorbic acid) was added in a ratio of 3:7 (v/v) and further incubated at 42°C for 30 mins. Absorbance was measured at 280 nm using a spectrophotometer. Standard graph was made with known amounts of KH_2_PO_4_ and total Pi content in the plant extracts calculated accordingly.

## Acknowledgements

All authors deeply acknowledge Regional Centre for Biotechnology (RCB), Faridabad for core-grant support, lab infrastructure, and central instrumental facilities. This study was supported by funds from DBT-Ramalingaswami Re-Entry Fellowship and Grant (No. BT/PR23666/AGIII/103/1039/2018) awarded to SB. The *phr1 phl1* double mutant seeds were a kind gift from Prof. Tzyy-Jen Chiou, Academia Sinica, Taipei. Prof. Ashis Kumar Nandi, Jawaharlal Nehru University (JNU), India provided the *npr1-1* seeds. D.L. acknowledges the Indian Institute of Science, India for start-up funds.

## Author Contributions

SB conceived the research. HG, KG, JJN, YW, and SB designed the experiments. HG, SB, KG, and KDI performed mutant identification via genotyping. HG, KG, KDI, YW, and AR performed pathogen growth assays and qPCRs. JJN, KDI, and YW performed protein detections analysis. DL and GS generated the InsP profiles in the mutants. YW and SB wrote the manuscript.

## Data availability statement

The data that supports the findings of this study are available in the supplementary material of this article

