## Supplementary Information for "Arabidopsis inositol polyphosphate kinases IPK1 and ITPK1 modulate crosstalks between SA-dependent immunity and phosphate-starvation responses"

**SUPPORTING INFORMATION**

Supporting information is available in the online version.

**FIGURE S1** Growth defects of *ipk1-1* are restored in the complemented lines *ipk1-1*:*Myc-IPK1*.

**FIGURE S2** In *ipk1-1* or *itpk1-2* complemented lines or their combinations with *phr1phl1* mutations, constitutive PSR and increased Pi-accumulations are abolished.

**FIGURE S3** The *vih2-4* plants display elevated total SA levels and upregulated expression of SA-responsive transcripts.

**FIGURE S4** Elevated PTI-responsive gene expressions in *ipk1-1* or *itpk1-2* are *SID2*-dependent.

**FIGURE S5** Upregulated defense-associated transcript levels, and constitutive PSI-gene expressions in *ipk1-1* are partially *NPR1*-dependent. Developmental defects however are *NPR1*-independent.

**FIGURE S6** SAX-HPLC profiles of $^{3}$H-labeled InsPs in Col-0, *itpk1-2*, *itpk4-1*, *mik-1*, and *ipk2β-1* plants.

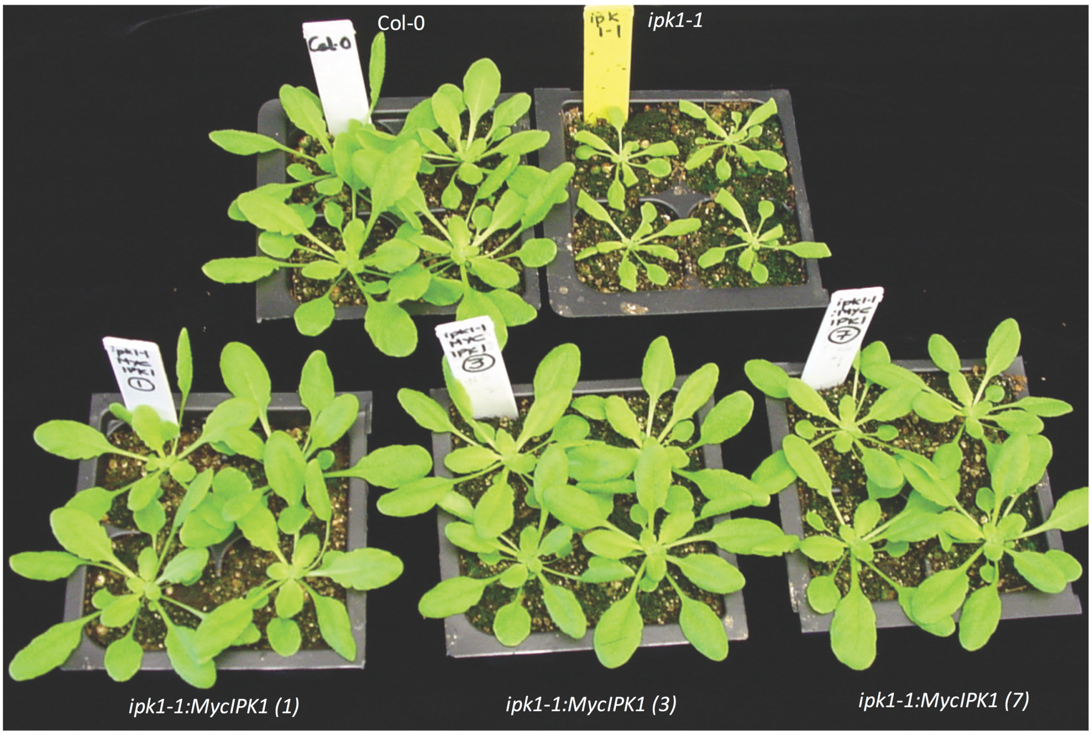


**FIGURE S1 Growth defects of *ipk1-1* are restored in the complemented lines *ipk1-1*:*Myc-IPK1*.** Plant Phenotypes of 4 week-old Col-0, *ipk1-1*, *ipk1-1:Myc-IPK1* Complemented lines #1, 3 and 7. The complemented line #1 was used for further assays.

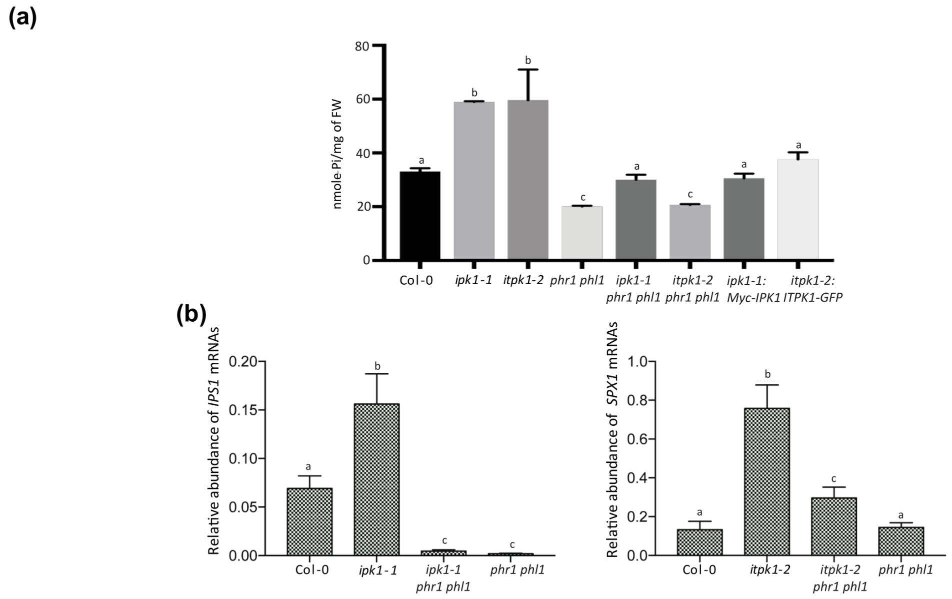


**FIGURE S2 In *ipk1-1* or *itpk1-2* complemented lines or their combinations with *phr1phl1* mutations, constitutive PSI-gene expressions and increased Pi-accumulations are abolished.** (a) Endogenous Pi levels in Col-0, *ipk1-1*, *itpk1-2*, *phr1 phl1*, *ipk1-1 phr1 phl1, itpk1-2 phr1 phl1*, *ipk1-1:Myc-IPK1*, and *itpk1-2:ITPK1-GFP*. (b) Relative expression levels of *IPS1* and *SPX1* in Col-0, *ipk1-1*, *itpk1-2*, *phr1 phl1*, *ipk1-1 phr1 phl1* or *itpk1-2 phr1 phl1,* respectively. The qRT-PCRs are normalized to the internal control *MON1* gene expression levels. All data presented here are mean values ± SD of three biological and technical replicates (n=3). Different letters mark statistically significant differences according to post-hoc Tukey's test (p<0.05).

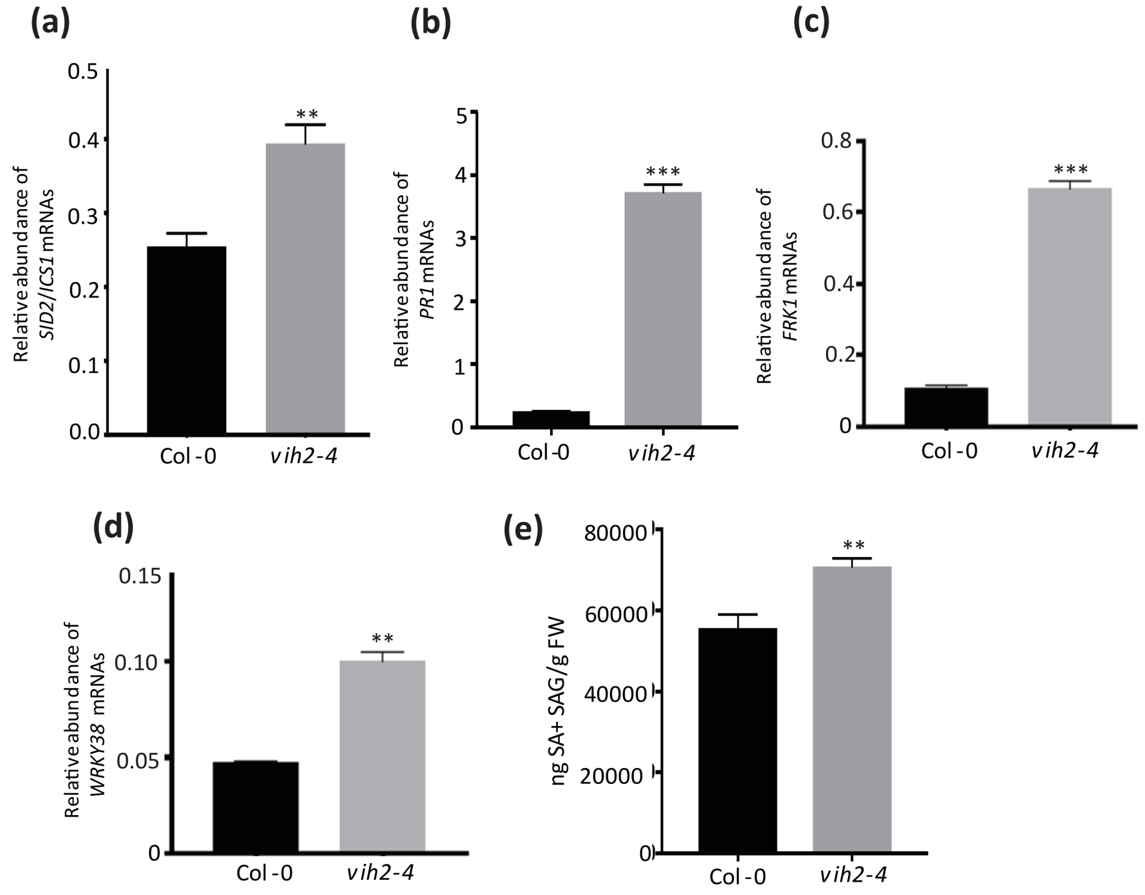


**FIGURE S3** ***vih2-4*** **plants display elevated total SA levels and upregulated expression of SA-responsive transcripts.** (a-d) Relative expression levels of *SID2/ICS1*, *PR1*, *FRK1*, and *WRKY38* in Col-0 and *vih2-4* plants. Expression of *MON1* gene was used as an internal control in the qRT-PCR assays. (e) Endogenous levels of total SA+SAG levels in Col-0 and *vih2-4*. SA estimations were performed on 3-4 week-old plants. Values indicate mean $\pm$ SD (three technical replicates, n=3). Student's *t*-test analysis was performed to calculate statistical significance (**p<0.001, ****p*<0.0001).

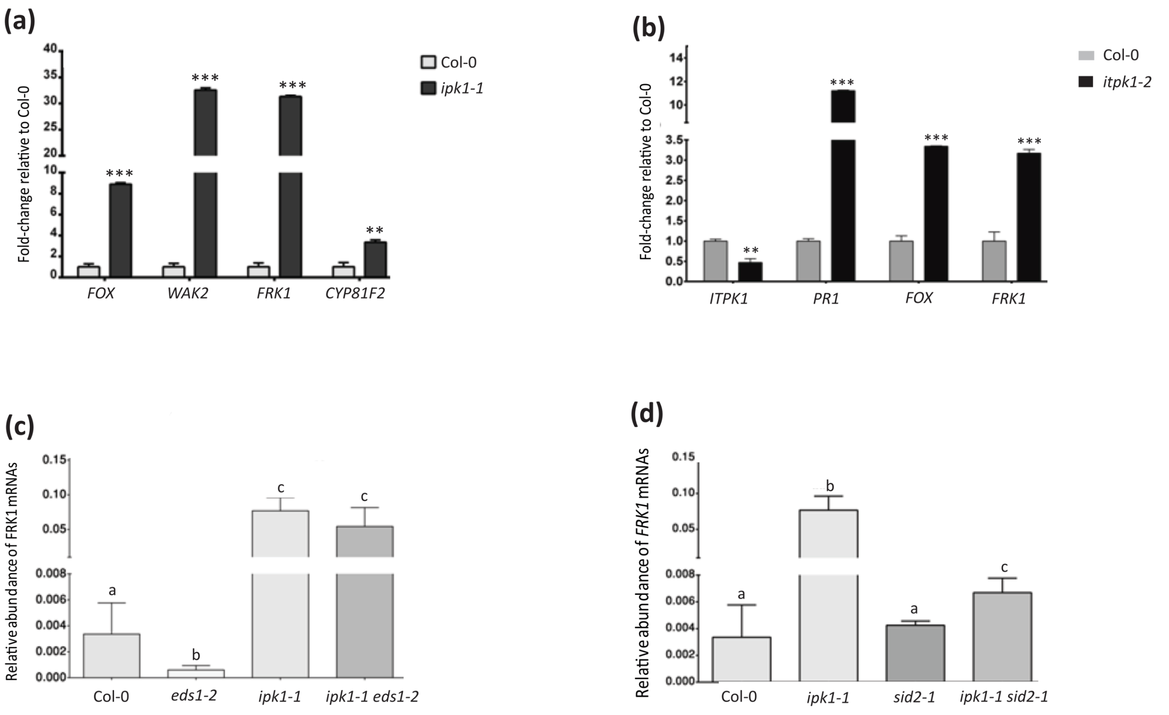


**FIGURE S4 Elevated PTI-responsive gene expressions in *ipk1-1* are *SID2*-dependent.** Fold-change in endogenous expression levels of basal defense-associated transcripts in (a) *ipk1-1*, or (b) *itpk1-2* plants compared to Col-0. Relative abundance of PTI marker *FRK1* in (c) *ipk1-1 eds1-2*, or (d) *ipk1-1 sid2-1* plants in comparison to Col-0, *ipk1-1*, *eds1-2*, or *sid2-1* plants. The qRT-PCRs data is relative to endogenous *MON1* gene expressions with mean ± SD of at least three biological and technical replicates (n=3). Statistical significance is with Student's *t*-test (**p<0.001, ***$p$<0.0001). Different alphabets are representative of post-hoc Tukey's test (p<0.05).

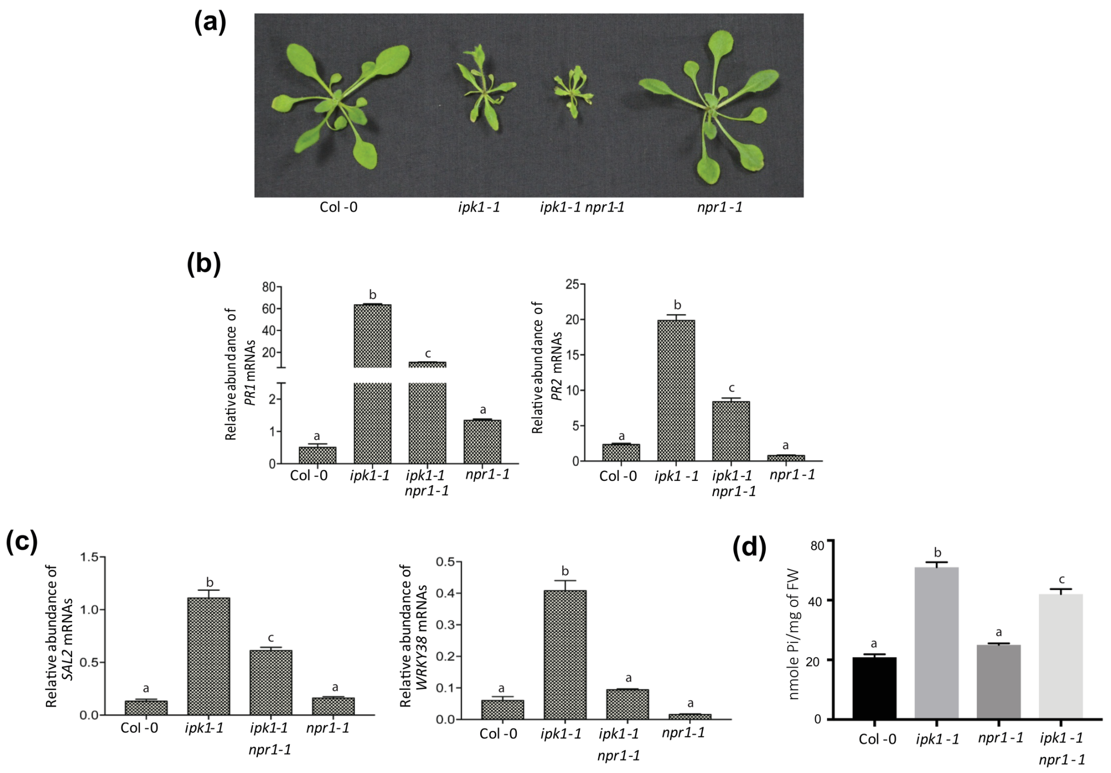


**FIGURE S5 Upregulated defense-associated transcript levels, constitutive PSI-gene expressions, or elevated basal Pi in *ipk1-1*, but not developmental defects, is partially *NPR1*-dependent.** (a) Growth phenotypes of 3-4 week-old Col-0, *ipk1-1*, *ipk1-1 npr1-1*, and *npr1-1* plants. (b-c) Relative abundance of *PR1*, *PR2*, *SAL2*, or *WRKY38* transcripts, respectively in indicated plants. The data is normalized to endogenous *MON1* gene expressions and are mean values (<u>+</u> SD) of 3 biological and technical replicates. (e) Endogenous Pi levels in indicated plants. Post-hoc Tukey's test ($p<0.05$) is shown by different letters.

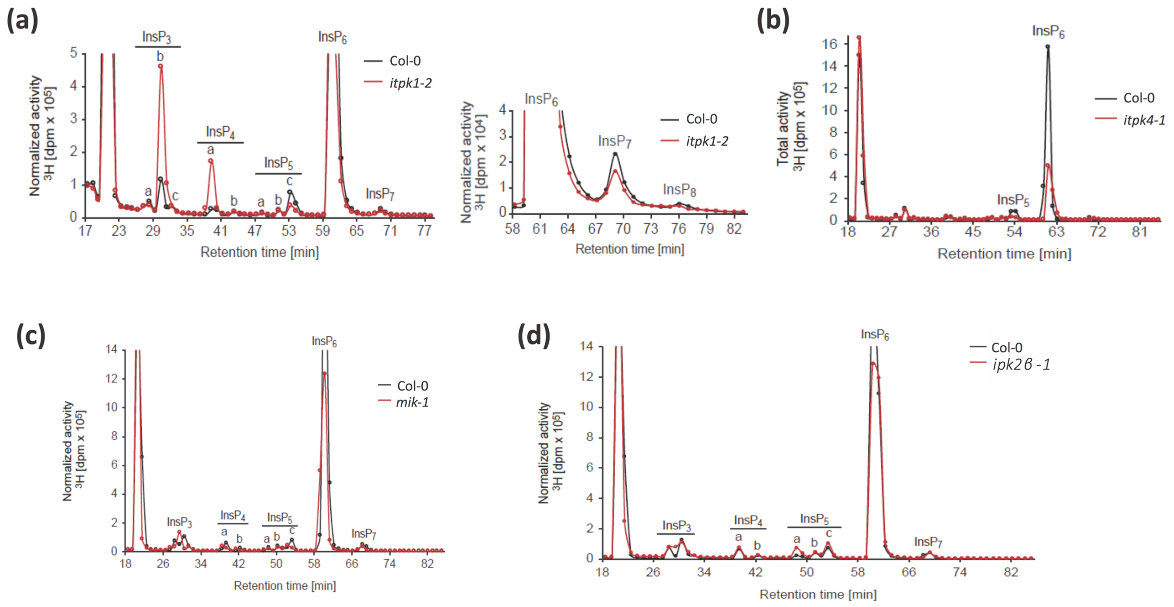


**FIGURE S6 SAX-HPLC profiles of $^3$H-labeled InsPs in Col-0, *itpk1-2*, *itpk4-1*, *mik-1*, and *ipk2β-1* plants.**

**SUPPLEMENTARY TABLE 1.** List of primers used.

| S. No | Primer Names | Primer Sequence (5’ to 3’) | Purpose (Ref) |
|---|---|---|---|
| 1. | *ipk1-1* RP | TTTCAATCAAGGAATGCTTGG | Genotyping *ipk1-1*<br>(Walia *et al*., 2020) |
| 2. | *ipk1-1* LP | ATGCTACGGGAAACCATTTTC | Genotyping *ipk1-1*<br>(Walia *et al.,* 2020) |
| 3. | *itpk1-2* RP | CCATGTCCCAGAAGAACTCAG | Genotyping *itpk1-2*<br>(Walia *et al*., 2020) |
| 4. | *itpk1-2* LP | ACCAATATTCGATTCGATTCCACACG | Genotyping *itpk1-2*<br>(Walia *et al*., 2020) |
| 5. | LB1.3 | ATTTTGCCGATTTCGGAAC | T-DNA Left-border primer (for SALK mutants)<br>(Walia *et al*., 2020) |
| 6. | LB3 | TAGCATCTGAATTTCATAACCAATCTC GATACAC | T-DNA Left-border primer (for SAIL mutants)<br>(Walia *et al*., 2020) |
| 7. | *ipk2α-1* RP | CACATTGCTAAAGACGGGAAG | Genotyping *ipk2α-1*<br>(This study) |
| 8. | *ipk2α-1* LP | ACCATTCTCAGTCAATCGTCG | Genotyping *ipk2α-1*<br>(This study) |
| 9. | *ipk2β-1* RP | TTGCTGGTCACATTGCTAGTG | Genotyping *ipk2β-1*<br>(This study) |
| 10. | *ipk2β-1* LP | TACCTGAAAACGCAATTTTGC | Genotyping *ipk2β-1*<br>(This study) |
| 11. | *itpk3-1* RP | TCGCCGTGTTCGTTAGTAATC | Genotyping *itpk3-1*<br>(This study) |
| 12. | *itpk3-1* LP | TGACTCACGACAAGGAAAGAG | Genotyping *itpk3-1*<br>(This study) |

| | | | |
|---|---|---|---|
| 13. | *mik-1* RP | CTTAGACGCTCATGGAACTCG | Genotyping *mik-1*<br>(This study) |
| 14. | *mik-1* LP | AAATGGTGGAGATCTGTGGACG | Genotyping *mik-1*<br>(This study) |
| 15. | *npr1-1* For | ATGTCTCGAATGTACATAAGGC | Genotyping *npr1-1*<br>(Ramirez *et al*., 2010) |
| 16. | *npr1-1* Rev | CTCAGTTTCCTAATAGAGAGG | Genotyping *npr1-1*<br>(Ramirez *et al*., 2010) |
| 17. | *itpk4-1* RP | TGTGAGAGCGAAATCAAAGTG | Genotyping *itpk4-1*<br>(This study) |
| 18. | *itpk4-1* LP | TTACATGCGTCAGCTGAGTTG | Genotyping *itpk4-1*<br>(This study) |
| 19. | *vih2-4* RP | AACAACAGCAATGACACAACG | Genotyping *vih2-4*<br>(Laha *et al*., 2015) |
| 20. | *vih2-4* LP | CATTCCCATCTTTTGGACAAC | Genotyping *vih2-4*<br>(Laha *et al*., 2015) |
| 21. | *eds1-2* For | CCCTTTCTAGTTTCCTTGAGCTAAG | Genotyping *eds1-2*<br>(Bhattacharjee *et al*., 2011) |
| 22. | *eds1-2* Rev | TCAGGTATCTGTTATTTCATCCATC | Genotyping *eds1-2*<br>(Bhattacharjee *et al*., 2011) |
| 23. | *phr1* RP | CTTTCTGGCGAACCTGTAGTG | Genotyping *phr1*<br>(This study) |
| 24. | *phr1* LP | GAGAGACCTCACACGCACTTC | Genotyping *phr1*<br>(This study) |
| 25. | *phl1* RP | TCCCACAATCCAAATTCAGAG | Genotyping *phl1*<br>(This study) |
| 26. | *phl1* LP | GTGGAGACGTTTCTGCACTTC | Genotyping *phl1*<br>(This study) |
| 27. | *sid2-1 RP* | AAGCAAAATGTTTGAGTCAGCA | Genotyping *sid2-1*<br>(Wildermuth *et al*., 2001) |
| 28. | *sid2-1* LP | TTCTTCATGCAGGGGAGGAG | Genotyping *sid2-1*<br>(Wildermuth *et al*., 2001) |

| | | | |
|---|---|---|---|
| 29. | qRT-PCR *PR1* For | GTGGGTTAGCGAGAAGGCTA | For qRT-PCRs<br>(Ingole *et al*., 2020) |
| 30. | qRT-PCR *PR1* Rev | ACCTTGGCACATCCGAGTCT | For qRT-PCRs<br>(Ingole *et al*., 2020) |
| 31. | qRT-PCR *PR2* For | TCAAGGAAGGTTCAGGGATG | For qRT-PCRs<br>(Ingole *et al*., 2020) |
| 32. | qRT-PCR *PR2* Rev | TTCACGAGCAAGGGAGATTG | For qRT-PCRs<br>(Ingole *et al*., 2020) |
| 33. | qRT-PCR *FRK1* For | CGGTCAGATTTCAACAGTTGTC | For qRT-PCRs<br>(Ingole *et al*., 2020) |
| 34. | qRT-PCR *FRK1* Rev | AATAGCAGGTTGGCCTGTAATC | For qRT-PCRs<br>(Ingole *et al*., 2020) |
| 35. | qRT-PCR *SPX1* For | GATTCCATTGTTGGAGCAAGA | For qRT-PCRs<br>(Walia *et al*., 2020) |
| 36. | qRT-PCR *SPX1* Rev | AATCTTGTTAGCTTCTTCTATTGTA | For qRT-PCRs<br>(Walia *et al*., 2020) |
| 37. | qRT-PCR *PHT3;2* For | GCTCCATCTGTAAGTGCATA | For qRT-PCRs<br>(This study) |
| 38. | qRT-PCR *PHT3;2* Rev | CTGGTAGTTCTGTTTCCCTT | For qRT-PCRs<br>(This study) |
| 39. | qRT-PCR MON1 For | AACTCTATGCAGCATTTGATCCACT | For qRT-PCRs<br>(Ingole *et al*., 2020) |
| 40. | qRT-PCR *MON1* Rev | TGATTGCATATCTTTATCGCCATC | For qRT-PCRs<br>(Ingole *et al*., 2020) |
| 41. | qRT-PCR NPR1 For | CTTGCGGAGAAGACGACACT | For qRT-PCRs<br>(This study) |
| 42. | qRT-PCR *NPR1* Rev | CACCGACGACGATGAGAGAG | For qRT-PCRs<br>(This study) |
| 43. | qRT-PCR SAL2 For | AGGAGCTAGCCGCTGCAAAGAAAG | For qRT-PCRs<br>(This study) |
| 44. | qRT-PCR SAL2 Rev | TCACTTCCATTCTTCCGCAGAT | For qRT-PCRs |

| | | | |
|---|---|---|---|
| | | | (This study) |
| 45. | qRT-PCR WRKY38 For | CGCCATGCGGTTGAAGAG | For qRT-PCRs (This study) |
| 46. | qRT-PCR WRKY38 Rev | TAACTTGAAAGCGGTCCACCA | For qRT-PCRs (This study) |
| 47. | qRT-PCR ICS1 For | TACTAACCAGTCCGAAAGACG | For qRT-PCRs (Ingole *et al.*, 2020) |
| 48. | qRT-PCR ICS1 Rev | GAGGCTTGACAACAACTCTGT | For qRT-PCRs (Ingole *et al.*, 2020) |
| 49. | qRT-PCR PAP17 For | TAGAGCCGGCTAAAAGCGAC | For qRT-PCRs (This study) |
| 50. | qRT-PCR PAP17 Rev | ATCTCCCGTAGACACCACGA | For qRT-PCRs (This study) |
| 51. | qRT-PCR IPS1 For | TCCCTCTAGAAATTGGGCAAC | For qRT-PCRs (Walia *et al.*, 2020) |
| 52. | qRT-PCR IPS1 Rev | GGGAGTGGGTACAACCCAAA | For qRT-PCRs (Walia *et al.*, 2020) |
